# Impaired hematopoiesis and leukemia development in mice with a conditional “knock-in” allele of a mutant splicing factor gene *U2af1*

**DOI:** 10.1101/345538

**Authors:** Dennis Liang Fei, Tao Zhen, Benjamin Durham, John Ferrarone, Tuo Zhang, Lisa Garrett, Akihide Yoshimi, Omar Abdel-Wahab, Robert K. Bradley, Paul Liu, Harold Varmus

**Affiliations:** Department of Medicine, Meyer Cancer Center, Weill Cornell Medicine, New York, NY 10065, United States; Cancer Biology Section, Cancer Genetics Branch, National Human Genome Research Institute, Bethesda, MD 20892, United States; Oncogenesis and Development Section, National Human Genome Research Institute, National Institutes of Health, Bethesda, MD 20892, United States; Human Oncology and Pathogenesis Program, Memorial Sloan Kettering Cancer Center, New York, NY 10065, United States; Genomics Resources Core Facility, Department of Microbiology and Immunology, Weill Cornell Medicine, New York, NY 10065, United States; Embryonic Stem Cell and Transgenic Mouse Core, National Human Genome Research Institute, National Institutes of Health, Bethesda, MD 20892, United States; Leukemia Service, Department of Medicine, Memorial Sloan Kettering Cancer Center, New York, NY 10065, United States; Computational Biology Program, Public Health Sciences Division, Fred Hutchinson Cancer Research Center, Seattle, WA 98109, United States; Basic Sciences Division, Fred Hutchinson Cancer Research Center, Seattle, Washington 98109, United States; New York Genome Center, New York, NY 10013, United States

**Keywords:** U2AF1, splicing factor, S34F, Runx1, myelodysplastic syndromes, leukemia

## Abstract

Mutations affecting the spliceosomal protein U2AF1 are commonly found in myelodysplastic syndromes (MDS) and secondary acute myeloid leukemia (sAML). We have generated mice that carry Cre-dependent “knock-in” alleles of *U2af1*(S34F), the murine version of the most common mutant allele of *U2AF1* encountered in human cancers. Cre-mediated recombination in murine hematopoietic lineages caused changes in RNA splicing, as well as multilineage cytopenia, macrocytic anemia, decreased hematopoietic stem and progenitor cells, low-grade dysplasias, and impaired transplantability, but without lifespan shortening or leukemia development. In an attempt to identify *U2af1*(S34F)-cooperating changes that promote leukemogenesis, we combined *U2af1*(S34F) with *Runx1* deficiency in mice and further treated the mice with a mutagen, N-Ethyl-N-Nitrosourea (ENU). Overall, three of sixteen ENU-treated compound transgenic mice developed AML. However, AML did not arise in mice with other genotypes or without ENU treatment. Sequencing DNA from the three AMLs revealed somatic mutations homologous to those considered to be drivers of human AML, including predicted loss-or gain-of-function mutations in *Tet2, Gata2, Idh1*, and *Ikzf1.* However, the engineered *U2af1(S34F)* missense mutation reverted to wild type (WT) in two of the three AML cases, implying that *U2af1*(S34F) is dispensable, or even selected against, once leukemia is established.

**SIGNIFICANCE STATEMENT:** Somatic mutations in four splicing factor genes (*U2AF1, SRSF2, SF3B1*, and *ZRSR2*) are found in MDS and MDS-related AML, blood cancers with few effective treatment options. However, the pathophysiological effects of these mutations remain poorly characterized, in part due to the paucity of disease-relevant models. Here, we report the establishment of mouse models to study the most common *U2AF1* mutation, *U2af1*(S34F). Production of the mutant protein specifically in the murine hematopoietic compartment disrupts hematopoiesis in ways resembling human MDS. We further identified deletion of the *Runx1* gene and other known oncogenic mutations as changes that might collaborate with *U2af1*(S34F) to give rise to frank AML in mice.

## INTRODUCTION

Myelodysplastic syndromes (MDS) are neoplastic diseases characterized principally by deficiencies of normal cells in myeloid, erythroid, and/or megakaryocytic lineages, mono- or multi-lineage dysplasia, clonal dominance of abnormal immature cells, and variable risks of developing secondary AML (1–3). In over half of MDS patients, the malignant clone carries a mutation in one of four genes (*U2AF1, SRSF2, SF3B1*, and *ZRSR2*) encoding factors critical for correct splicing of pre-messenger RNA (pre-mRNA) (4). Mutations in these “splicing factor genes” are presumed to be causative events in MDS because of their frequency, their recurrence at a few positions in coding sequences, their high allelic ratios, and their association with clinical outcomes (4–7). Still, despite intensive studies, the mechanisms by which such mutations contribute to the initiation or maintenance of neoplasias have not been identified.

We have been studying the most common mutation observed in *U2AF1*, the gene encoding an RNA-binding protein that helps to direct the U2 small nuclear ribonucleoprotein particle (U2 snRNP) to the 3’ splice acceptor site in pre-mRNA (8–10). This mutation replaces serine at position 34 of U2AF1 with phenylalanine (S34F) (4, 6), yielding a neomorphic splicing factor that changes splicing patterns for many RNAs and does not preserve enough normal splicing to permit cell survival in the absence of a wild-type (WT) allele (11). In addition, *U2AF1* (S34F) is present at high allelic frequencies (near 50%) in MDS (6), seems to predispose MDS patients to secondary AML (6, 12), and is also found, albeit at lower frequencies, in a variety of other neoplastic diseases, including solid tumors (13–15).

Progress towards understanding the oncogenic effects of *U2af1*(S34F) has been impeded by a lack of appropriate biological models. Here we describe our development of genetically engineered mouse models in which *U2af1*(S34F) is assembled at the endogenous *U2af1* locus by the action of the Cre recombinase. By activating Cre in the hematopoietic compartment of these mice to produce *U2af1*(S34F), the mice develop impairments of blood cells with MDS-like features, accompanied by abnormal splicing patterns resembling those previously observed in human cells expressing this mutant splicing factor. In an effort to model *U2AF1* (S34F)-associated leukemia, we deprived the *U2af1* mutant mice of a hematopoietic transcription factor, Runx1, often co-mutated in human MDS and leukemias (16, 17), and treated them with a chemical mutagen, N-Ethyl-N-Nitrosourea (ENU). Under those circumstances, clones of AML developed in three of sixteen mice, but not in mice lacking any of the three factors. Although AML did not occur often enough to conclude that the mutant splicing factor is required for leukemogenesis in this model, whole exome sequencing of AML cells revealed somatic mutations in genes often mutated in human AML. In summary, we report the first “knock-in” mouse lines with conditional *U2af1*(S34F) alleles, demonstrate the effects of mutant U2af1 on mouse hematopoiesis, and identify additional mutations that may cooperate with *U2af1*(S34F) and *Runx1* deficiency during leukemogenesis.

## RESULTS

### Establishing mice carrying conditional knock-in S34F alleles of *U2af1*

We used targeting vectors (termed *MGS34F* and *IES34F*; Figs 1A and S2A) to create two conditional S34F mutant alleles at the endogenous *U2af1* locus of B6/129 mice, and we documented the successful introduction of the targeting vectors at the appropriate sites by restriction mapping and Sanger sequencing (Figs S1A, S2B, and Methods). Mice carrying either of the altered *U2af1* alleles express green fluorescent protein (GFP) in all tissues because the mouse embryonic stem cell line used to generate the mutant mice carries a *GFP* transgene driven by the ubiquitously active human UBC promoter (18) and the transgene is located on the same chromosome as the *U2af1* locus, with a crossover frequency of 1.95% (Fig. S1H; Table S1).

**Figure 1.**
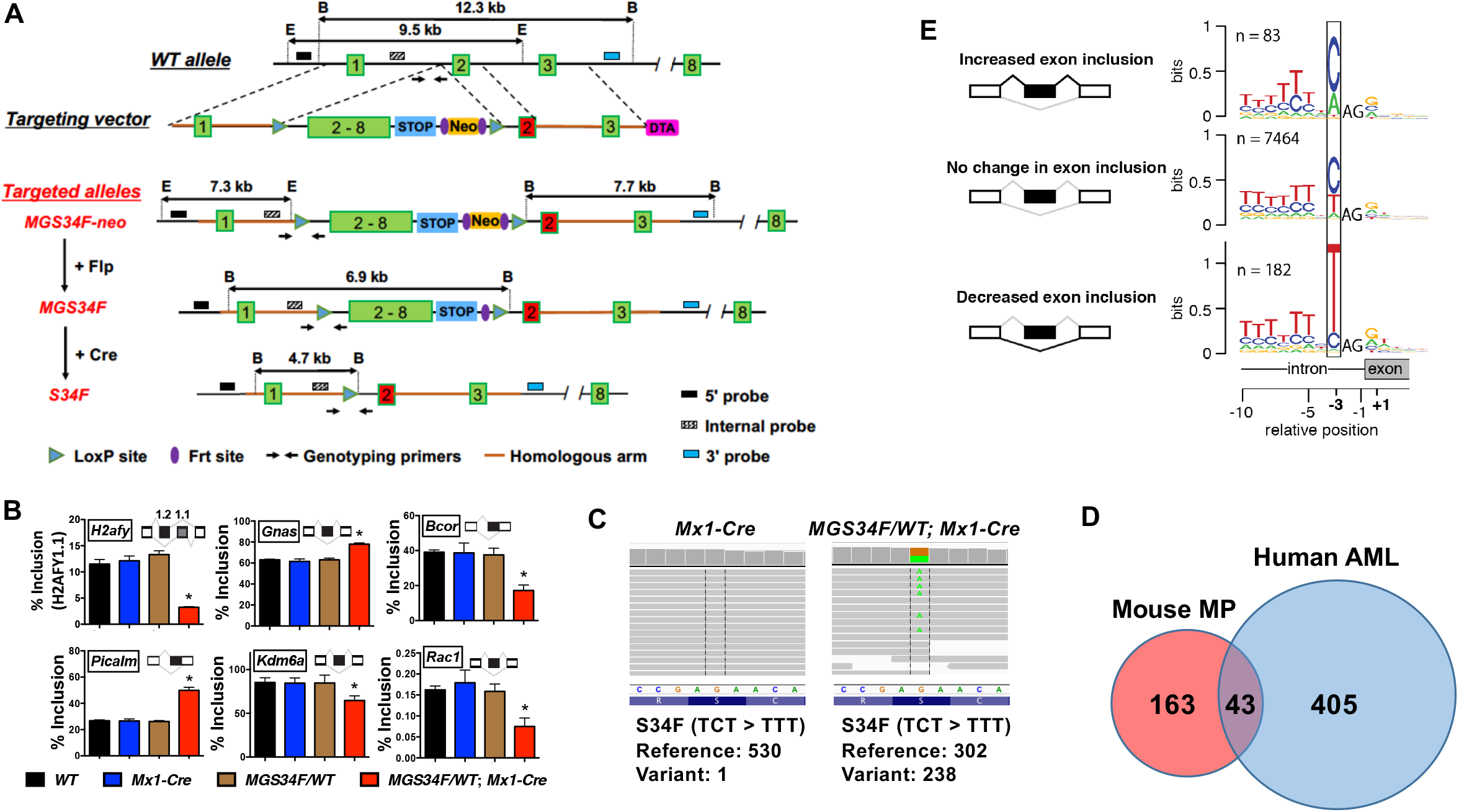
Conditional expression of *U2af1*(S34F) from the mouse endogenous locus alters splicing similar to those in human cells expressing *U2af1*(S34F). (**A**) Diagrams of the endogenous *U2af1* locus, the *MGS34F* targeting vector, and modified alleles, with sites used for Southern- and PCR-based genotyping. Numbers in boxes indicate exons or exonic sequences in cDNA; lines represent introns; STOP denotes a 3x transcriptional stop signal from SV40; E, EcoRI site; B, BamHI site. The red version of Exon 2 encodes the S34F mutation (TCT > TTT). A more detailed description of the targeting vector is in the Supplemental Method section. (**B**) *U2af1*(S34F) changes the inclusion levels of indicated exons (or portions of exons, see the inserted cartoons) found at six loci of genes (named in boxes) in total bone marrow cells from mice with the indicated genotypes (n = 3). The mice were treated with poly (IC), then euthanized two weeks later for RNA extraction. As noted in the text, alternative splicing of human homologs of these mRNAs was previously reported to be affected by *U2af1*(S34F). Asterisks indicate statistically significant changes compared to other genotypes by t test (p < 0.05). Error bars represent standard error of the mean (s.e.m). (**C**) Representative read coverage of *U2af1* cDNA by RNA-seq, which was performed on MPs from mice of indicated genotypes four weeks after poly (IC) treatment. Each grey line represents a sequencing read. The reference DNA (non-coding strand) and protein sequences are shown below the sequencing reads, and the numbers of reference (WT) and variant (S34F) alleles are quantified (bottom of the graphs). (**D**) The Venn diagram indicates the numbers of orthologous genes from mouse and human datasets that show at least 10% change in cassette exon inclusion levels in the presence of mutant U2AF1. The mouse dataset is presented in Supplemental Table S3. The human dataset was published previously (21), using AML samples from TCGA (61). Genes with low levels of expression (median transcripts per million sequenced RNAs < 10) or have no mouse or human ortholog were excluded from the analysis. (**E**) *U2af1*(S34F) recognizes similar consensus sequences at 3’ splice sites in mouse genome as *U2af1*(S34F) does in human genome. mRNA from MPs with or without *U2af1*(S34F) was sequenced to determine nucleotides at the 3’ splice acceptor sites of cassette exons and displayed as sequence logos according to whether inclusion of the exon in mRNA was increased, decreased, or unaffected by *U2af1*(S34F). The resemblance of these logos to those previously determined with human materials is discussed in the text. Additional molecular characterization of the established mouse lines (including another conditional allele, IES34F), and their cell derivatives are presented in Supplemental Figures S1 and S2.

To assess whether these conditional alleles can be rearranged correctly, we crossed the two mouse strains with mice carrying a UBC-driven transgene encoding a tamoxifen-dependent Cre recombinase (UBC-CreERT2) (19). Mouse embryo fibroblasts (MEFs) derived from embryos with *MGS34F; CreERT2* or *IES34F; CreERT2* were treated with 4-hydroxyl-tamoxifen (4OHT) in culture; efficient appearance of the expected configurations of *U2af1*(S34F) was confirmed by Southern blotting (Figs S1B and S2C), and the anticipated changes in *U2af1*(S34F) mRNA were ascertained with allele-specific Taqman assays (Figs S1C, S2D, and S3C – E). *MGS34F* appeared to be more efficiently recombined by Cre than was the *IES34S* allele, and was used in all follow-up studies.

We next crossed mice heterozygous for the conditional *MGS34F* allele with mice carrying an *Mx1-Cre* transgene that is expressed in the blood lineage upon administration of poly (IC) (20). Two weeks after poly (IC) treatment of the resulting bi-transgenic mice (*MGS34F/WT; Mx1-Cre*), Cre-mediated recombination was nearly complete in nucleated bone marrow cells, as revealed by Southern blotting (Fig. S1E) and by quantitative analysis of PCR-generated fragments (Fig. S3A and S3B). The WT and mutant alleles appear to be expressed at equivalent levels, judging from the approximately 1:1 ratio of mutant and WT *U2af1* mRNA (Fig. S1F).

### Effects of *U2af1*(S34F) on pre-mRNA splicing in mouse cells resemble effects of *U2af1*(S34F) in human cells

*U2AF1*(S34F) is known to affect pre-mRNA splicing and possibly other events in a variety of human cells (11, 21–27). We used cells and tissues from mice carrying the *U2af1(S34F)* allele to determine whether similar alterations occur in mice after expression of *U2af1*(S34F) at physiological levels. As an initial assessment, we measured changes in spliced products of pre-mRNA from six mouse genes (*H2afy, Gnas, Bcor, Picalm, Kdm6a*, and *Rac1*) homologous to human genes whose transcripts undergo changes in alternative splicing (11, 21, 27) in the presence of *U2af1*(S34F). We also examined RNA products of one mouse gene, *Atg7*, whose transcripts were reported to show changes in use of polyadenylation sites (25). Isoform sensitive primers were designed to target regions of the mRNAs that are predicted to undergo changes based on previous results in human or mouse cells (Table S2). We found that similar alterations in pre-mRNA splicing occurred in the six homologous murine genes expressed in total bone marrow cells (Fig. 1B). However, the previously reported changes in *Atg7* RNA polyadenylation sites were not observed in mouse bone marrow cells expressing *U2af1*(S34F) (Fig. S1G).

We used whole cell mRNA sequencing methods to make a more extensive survey of *U2af1*(S34F)-induced alterations in pre-mRNA splicing, looking specifically for the changes in frequency with which so-called “cassette exons” are sometimes included in the spliced products, since these are the most prominent consequences of the S34F mutant in human cells (11, 21–24, 27). In addition, we performed these analyses in mouse myeloid progenitors (MPs), which were isolated from the bone marrow of *MG(S34F)/WT; Mx1-Cre* mice (and from *Mx1-Cre* mice as a control) four weeks after poly (IC) treatment based on their immunophenotype, Lin”Sca-1”c-Kit^+^ (Lin, Lineage), and expressed *U2af1*(S34F) mRNA as expected (Fig. 1C).

We found significant changes in the alternative splicing of cassette exons from 206 mouse genes; 43 of those genes (about 20 percent) are homologs of human genes similarly affected by *U2af1*(S34F) or U2AF1(S34Y) in AML cells (Fig. 1D). Furthermore, the consensus nucleotide sequences preceding the invariant AG dinucleotide at the 5’ end of the affected cassette exons match those previously identified as determinants of altered splicing patterns in human cells expressing *U2af1*(S34F) (11, 21–24, 27). For example, the first nucleotide position 5’ of the AG dinucleotide is frequently occupied with a C or A nucleotide upstream of exons that are more frequently included in mRNA, and is often occupied with a T nucleotide upstream of those cassette exons that are less frequently included (Fig. 1E). Similar changes were seen in MEFs expressing *U2af1*(S34F) (Figs S1D and S2E). Based on these results, we conclude that physiological expression of *U2af1*(S34F) causes changes in pre-mRNA splicing similar but not identical to those observed by us and others in human cells expressing *U2af1*(S34F).

### U2af1(S34F) affects hematopoiesis in mice, with features that are shared with human MDS

We next examined the impact of *U2af1*(S34F) on hematopoiesis in our genetically engineered mice. Blood samples were taken before and after administration of poly (IC) to induce Cre and subjected to complete blood cell counts (CBCs) and flow cytometry. The cellular profiles of blood from WT mice and from mice carrying only the *Mx1-Cre* transgene or only the *MGS34F* allele were similar and normal, as expected. However, *MGS34F/WT; Mx1-Cre* mice showed mild but persistent changes in red blood cells (reduced RBC count, hemoglobin concentration, and hematocrit; and increased mean red cell volume; Figs 2A, 2B, S4A, and S4B). Platelet numbers were normal (Fig. S4C), but the numbers of white blood cells were reduced persistently by nearly 50% compared to results with mice with the control genotypes after poly (IC) treatment (Fig. 2C). Decreases were observed in all major WBC lineages as measured by flow cytometry (Fig. 2D – 2G), with the most marked reduction seen in B220+ B cells (Fig. S4D – S4G).

**Figure 2.**
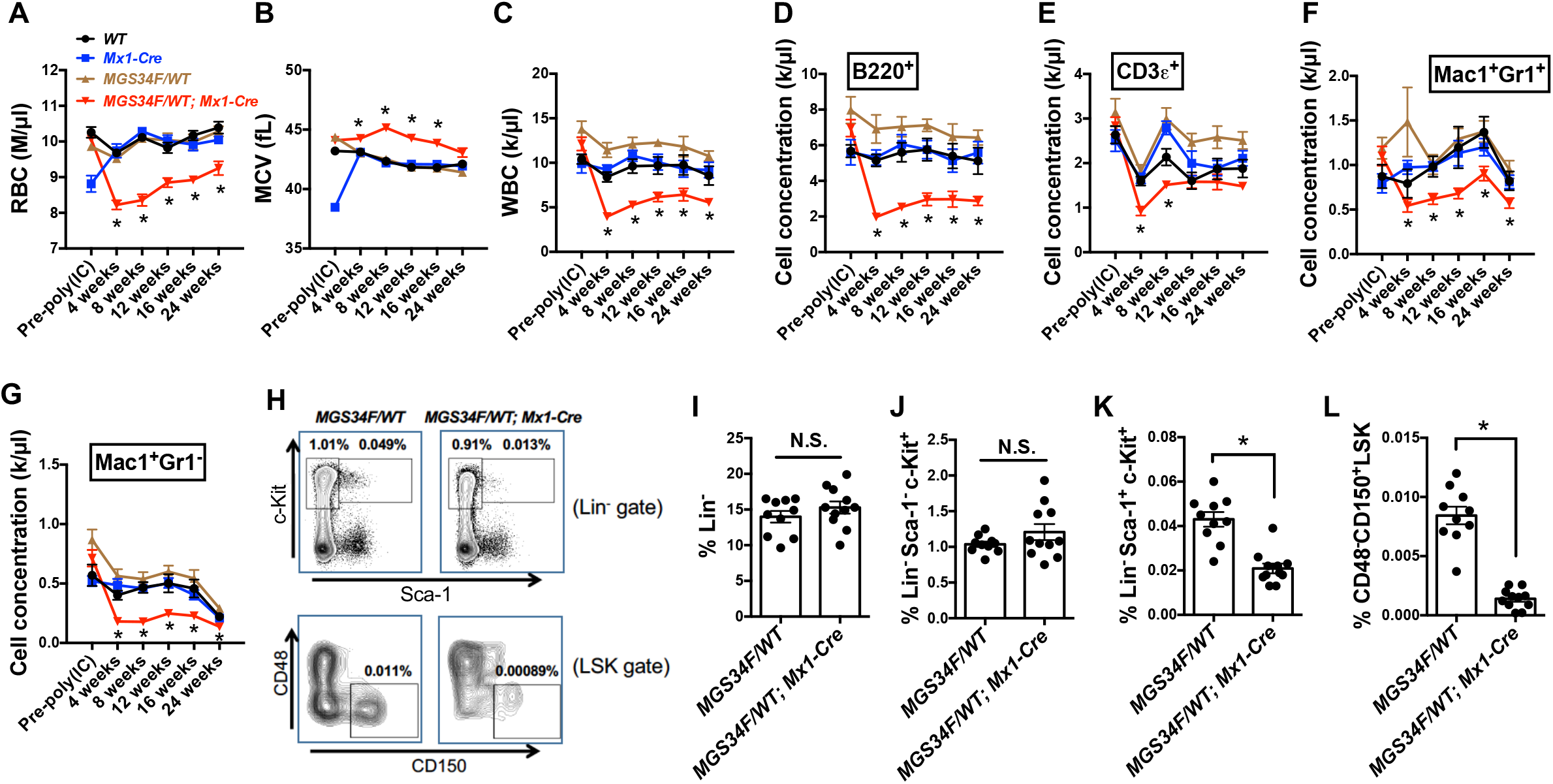
*U2af1*(S34F) causes multi-lineage cytopenia, macrocytic anemia, and reduction in the hematopoietic stem and progenitor cell populations. (**A – G**) Multilineage cytopenia and macrocytic anemia in mice expressing *U2af1*(S34F). Blood samples taken from mice of the indicated genotypes (see keys in panel **A**) before and up to 24 weeks after poly (IC) treatment were subjected to complete blood count to determine numbers of red blood cells (**A**), mean red blood cell volume (**B**), and white blood cells (**C**); and used for flow cytometry (panels **D – G**) to determine concentrations of cells with indicated lineage markers. Asterisks indicate significant changes for mice with the *MGS34F/WT; Mx1-Cre* genotype as compared to mice of any other genotype by multiple t test (FDR < 0.05). (**H – L**) *U2af1*(S34F) reduces the percentage of hematopoietic stem and progenitor cells in the bone marrow. Bone marrow cells from mice of the indicated genotypes were harvested 36 weeks after poly (IC) treatment, and the percentages of hematopoietic stem and progenitor cells were determined by flow cytometry using the stated markers. (**H**) Representative results of flow cytometry with numbers indicating the percentages of live nucleated cells in boxed areas. (**I – L**) Bar heights indicate the mean percentages of cells with the designated phenotypes (vertical axes) in the bone marrow. A dot indicated the percentage of cells in a mouse. Asterisks denote significant changes by t test (p < 0.05). N.S. not significant. Error bars represent s.e.m. See **Supplemental Figure S4** for additional characterization of these mice. *WT/WT* (n = 10), *WT/WT; Mx1-Cre* (n = 10), *MGS34F/WT* (n = 10), *MGS34F/WT; Mx1-Cre* (n = 11).

Despite the reduction in nearly all mature blood cell types upon activation of *U2af1*(S34F), bone marrow cellularity and spleen size were normal (Fig. S4I and S4J), with no evidence of pathological extramedullary hematopoiesis in mice with *U2af1*(S34F) (data not shown).

We sought to determine whether these long-term changes were due to effects of the mutant splicing factor on blood stem and progenitor cells. We examined the abundance and function of hematopoietic stem cells (HSCs) and progenitors in mutant and control mice 36 weeks after poly (IC) treatment. Bone marrow cells from mice with or without *U2af1*(S34F) were stained with antibodies against markers for HSCs and for various blood cell progenitors and were analyzed by flow cytometry. While no differences were observed in the percentages of Lin^-^ cells (Fig. 2I) and MPs (Lin^-^Sca-1^-^c-Kit^+^, Fig. 2H and 2J) from mutant and control animals, the LSK cells (marked by Lin^-^Sca-1^+^c-Kit^+^) and, especially, HSC-enriched cell population (marked by CD48^-^D150^+^ within the LSK cells) were decreased in the bone marrow of *U2af1* (S34F) mice (Figs 2H, 2K and 2L). These findings suggest that reductions in HSCs likely account for the persistence of multilineage cytopenia observed in the peripheral blood of mice expressing mutant U2af1.

To determine whether the effects of *U2af1* (S34F) on mouse hematopoiesis were cell autonomous, we transplanted bone marrow cells from *MGS34F/WT; Mx1-Cre* mice and from control *(MGS34F/WT)* mice non-competitively into lethally irradiated recipient mice (F1 progeny of a B6 × 129 cross; see Methods). Four weeks after transplantation, poly (IC) was used to induce production of Cre in the transplanted cells. Five weeks later, macrocytic anemia, multi-lineage cytopenia, and reduction in LSK cells were observed in the recipient mice (compare results in Fig. S5 with those in Figs 2 and S4). Although the numbers of LSK cells were reduced, a slightly higher fraction of these cells were actively cycling, as shown by a BrdU incorporation assay (Fig. S5Q), suggesting a compensatory mechanism to overcome the loss of these cells. Moreover, bone marrow cells from the *U2af1*(S34F) mice formed fewer myeloid colonies, especially after replating (Fig. S5R), consistent with the reduced numbers of LSK cells observed by flow cytometry. Overall, these results demonstrate that the effects of *U2af1*(S34F) on hematopoiesis are cell-autonomous.

To further characterize the HSC defects attributed to *U2af1*(S34F), we performed competitive transplantations using bone marrow cells from *MGS34F/WT; Mx1-Cre* or *MGS34F/WT* mice (test cells) mixed with bone marrow cells from WT or *Mx1-Cre* mice (competitor cells). Equal numbers of test and competitor cells were mixed and transplanted into lethally irradiated recipient mice, and the recipients received poly (IC) four weeks later (Fig. 3A). Mice that received test cells from *MGS34F/WT; Mx1-Cre* animals displayed a deficiency of U2af1(S34)-expressing cells in all measured mature and immature cell lineages in the peripheral blood, bone marrow, and spleen of the recipient mice, as early as two weeks after poly (IC) (Fig. 3B – 3F). In contrast, immature and mature blood cells from *MGS34F/WT* mice remained at levels of nearly 50% at all times. These results show that HSCs expressing *U2af1*(S34F) are defective in repopulating the hematopoietic compartment in a competitive transplant setting.

**Figure 3.**
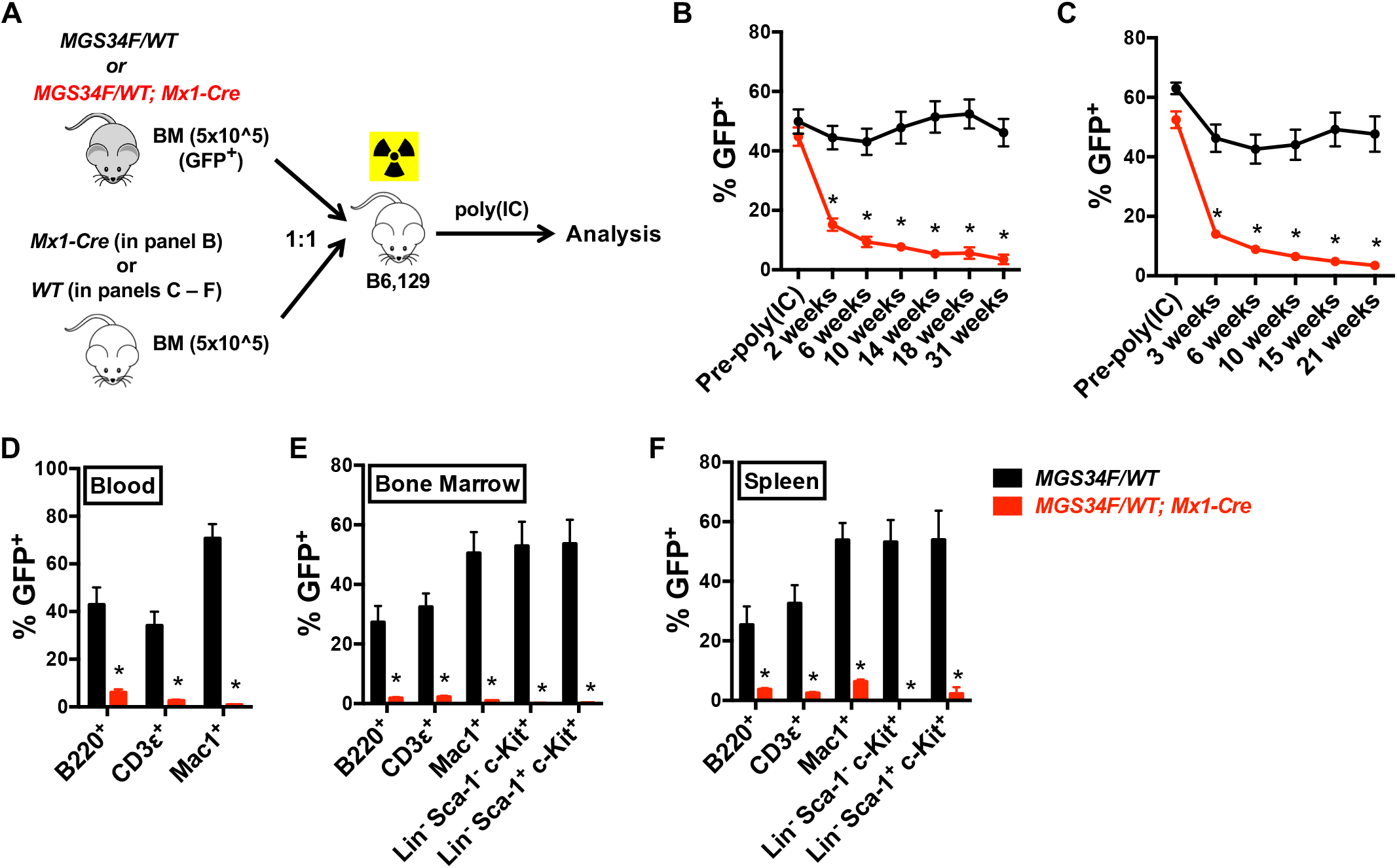
Hematopoietic stem and progenitor cells with *U2af1*(S34F) are eliminated in a competitive transplant assay with cells lacking mutant U2af1. (**A**) Assay design. Equal numbers (0.5 million nucleated cells) of donor *(MGS34F/WT* or *MGS34F/WT; Mx1-Cre*, both GFP^+^) and competitor (WT or *Mx1-Cre* alone, GFP”) bone marrow cells were mixed and transplanted to lethally irradiated, WT recipient mice. Recipient mice were treated with poly (IC) four weeks after transplant. (**B – F**) Contributions of donor cells to hematopoietic compartments of recipient mice. (**B** and **C**) *U2af1*(S34F)-containing BM cells from *MGS34F/WT; Mx1-Cre* mice, but not BM cells from *MGS34F/WT* mice, constituted less of the white blood cells with BM competitors from *Mx1-Cre* (**B**) or wildtype mice (**C**) as measured by percentages of GFP^+^ cells in peripheral blood in the recipient mice in a competitive transplant. (**D – F**) All measured cell lineages showed competitive disadvantages of *U2af1*(S34F)-containing cells in the indicated cell lineages in the blood (**D**), bone marrow (**E**), and spleen (**F**) 21 weeks after poly (IC) treatment. n = 5 −10 for each group of mice. Asterisks indicate significant changes by t test. Error bars represent s.e.m.

Using light microscopy to seek the morphological features of myelodysplasia, we occasionally observed bi-nucleated erythroid cells and hypo-segmented neutrophils in the bone marrow cells from *U2af1*(S34F)-expressing mice (Fig. S6A and S6B). Moreover, megakaryocytes from these mice occasionally formed clusters (Fig. S6C – S6E), even though platelet production was normal or only slightly diminished (Fig. S4C and S5C). These dysplastic features typically affected less than 1% of bone marrow cells, not present in the peripheral blood, and, therefore, did not meet the diagnostic criteria for human MDS. Nevertheless, the multi-lineage cytopenia and dysplasia found in the *U2af1*(S34F) mutant mice are hallmarks of human MDS.

### Generation of mice with conditional alleles of both *Runx1* and *U2af1*

Although *U2af1*(S34F) affects hematopoiesis in our genetically engineered mice, these mice were relatively healthy (e.g. with normal weight gain; Figs S4H and S5M), and they lived a normal life span. We reasoned that additional genetic changes might produce more severe hematologic defects or neoplasias dependent on *U2af1(S34F).* Therefore, we combined our *MGS34F* allele and the *Mx1-Cre* transgene with a *Runx1* allele carrying LoxP sites that flank exon 4, encoding the DNA-binding Runt domain (28). When bred to homozygosity at the floxed *Runx1* locus (*Runx1*^F/F^) in the presence of *Mx1-Cre* and *MGS34F*, Cre recombinase inactivates *Runx1* and allows production of the mutant splicing factor in the same cells after administration of poly (IC). Since loss-of-function mutations of *Runx1* often co-occur with *U2AF1* mutations in human myeloid neoplasms (16, 17), this seemed to be a promising genetic combination for eliciting pathological phenotypes, including leukemias.

The experimental cohort used for these purposes consisted of the “compound engineered” mice *(MGS34F/WT; Runx1*^FIF^; Mx1-Cre, called “URC mice”); mice in which poly (IC) activates *U2af1(S34F)* or inactivates *Runx1 (MGS34F/WT; Mx1-Cre* or *Runx1^F/F^; Mx1-Cre*); and control WT mice (or WT-equivalent mice lacking the *Mx1-Cre* transgene). After induction of Cre, URC mice developed the hematological abnormalities associated with either *U2af1*(S34F) (e.g. multi-lineage cytopenia, low-level myeloid and erythroid dysplasia) or *Runx1* deficiency alone (e.g. thrombocytopenia, increased percentage of myeloid cells, and increased myeloid colony formation; (28, 29)), as shown in Fig. S7A – S7D. We also examined the effects of *U2af1*(S34F) and *Runx1* deletion on gene expression and pre-mRNA splicing in MP cells from poly (IC)-treated URC mice and compared the findings with those obtained with mice with conditional alleles at either the *U2af1* or *Runx1* loci (Fig. S7E – S7G, Tables S3 and S4). As anticipated from its role as a gene encoding a transcription factor, *Runx1* deficiency affected mainly gene expression, whereas *U2af1*(S34F) changed predominantly mRNA splicing. As a result, MP cells with *U2af1*(S34F) and without functional *Runx1* exhibited many changes in both mRNA splicing and gene expression, as illustrated in Fig. S7F and S7G.

### URC mice develop frank AML or AML clones after ENU mutagenesis

Although URC mice showed many changes in hematopoiesis and in gene expression in MP cells (Fig. S7), they had a normal life span and did not develop frank leukemia within one and a half years after poly (IC) treatment. To seek additional genetic alterations that might cooperate with *U2af1*(S34F) and *Runx1* deficiency to cause more profound pathology, including leukemias, we performed a forward genetic screen by treating URC mice (6-16 weeks old) with the mutagen ENU one week after inducing programmed genetic changes with poly (IC) (Fig. 4A). We used a relatively low-dose of ENU (100 mg/kg) that, based on our previous experience (30), did not produce myeloid leukemia in otherwise normal animals. Mice that developed T cell malignancies and thymoma (known consequences of ENU) were excluded from further analysis.

**Figure 4.**
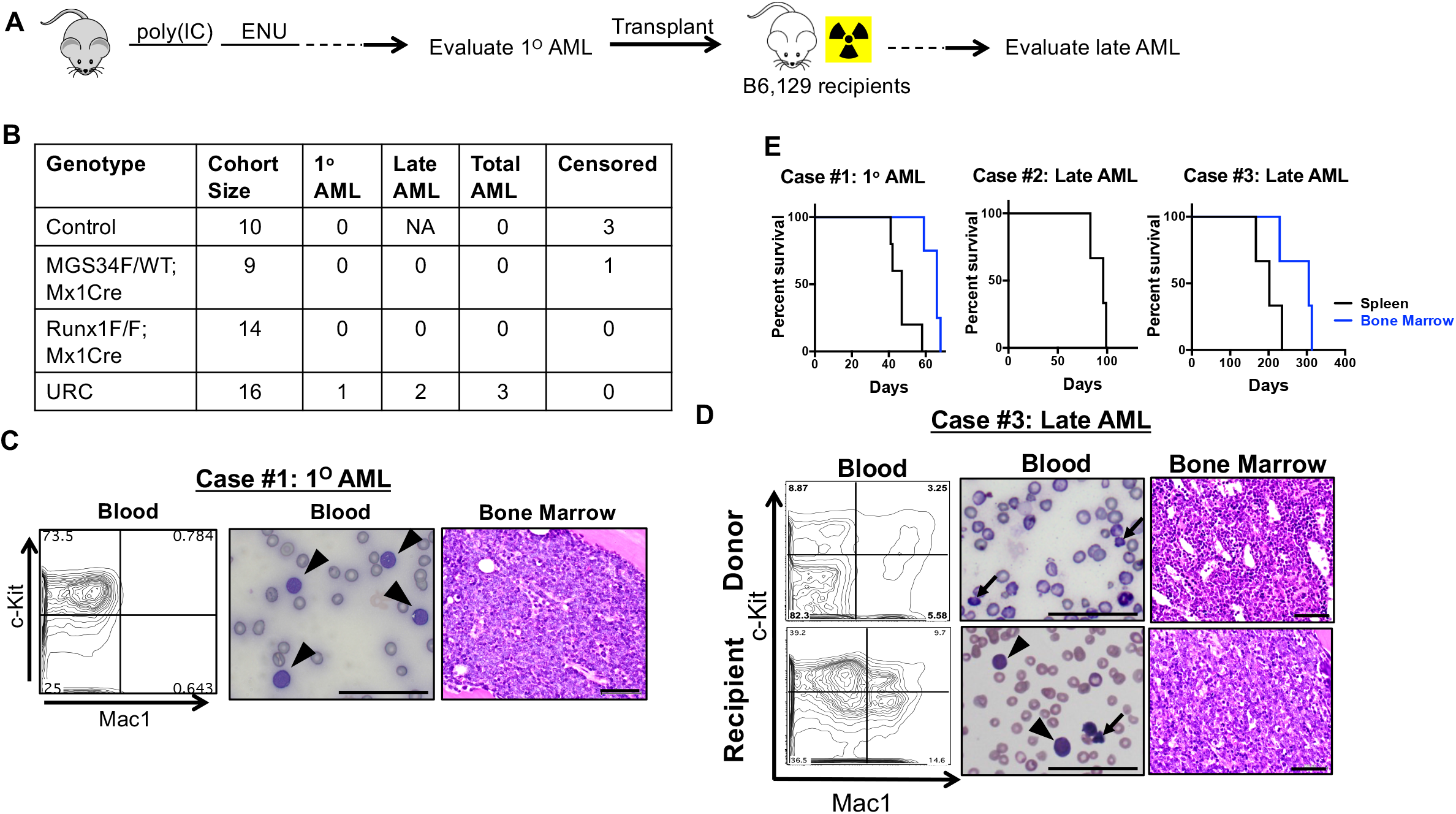
Leukemogenesis in ENU-treated URC mice. (**A**) Strategy for testing leukemogenesis. Mice were treated with poly (IC) and, one week later, with ENU (100 mg/kg), and then monitored until they were moribund or up to 1.5 years after receiving poly (IC). Splenic or bone marrow cells from these mice without frank AML were transplanted to sub-lethally irradiated, WT recipient mice. Three to five recipient mice were used for each transplant and monitored for up to one year for the development of AML. (**B**) Summary of AML incidences. Mice with the “control” genotype lack both *U2af1*(S34F) and *Runx1* deletions, and were not used for transplant. NA, not available. Mice with T cell lymphoma or thymoma were censored from the study cohort. In total, three of sixteen URC mice developed AML (p-value = 0.13, based on four separate genotypes as shown; p-value = 0.03 based on URC mice vs. all other genotypes, by Fisher’s exact test). (**C**) A case of primary AML in one of sixteen poly (IC)- and ENU-treated URC mice (Case #1). High percentage of c-Kit^+^ cells was present in the peripheral blood of this mouse by flow cytometry, which coincided with infiltration of morphologically defined myeloblasts (arrow head) in a Wright-Giemsa stained blood smear, and H&E stained section of the bone marrow. (D) A case of late AML. Representative results are shown for the Case #3 donor (top row) and one of transplant recipients when it was moribund (bottom row). Percentages of c-Kit^+^ cells were markedly increased in the peripheral blood of the recipient mouse, compared to that in the donor. Abnormally nucleated red blood cells (arrow) were observed in the blood smear from both the donor and recipient, while myeloblasts (arrow head) were only observed in blood from the recipient. Blast cells also filled the bone marrow of the recipient mouse. Scale bar = 50 μm. (**E**) Survival curves of sub-lethally irradiated mice after transplants of bone marrow (black) or spleen (blue) cells from the primary donor mice of Case #1 - #3. Day count started since after the transplantation. See Fig. S9 for additional characterization of these AML cases.

All mice lacking *Runx1* died prematurely (within 15 months after poly (IC) treatment), regardless of *U2af1* status, after receiving ENU (Fig. S8A). However, ten months after administration of ENU, one of the 16 URC mice (Case #1) — but none of the mice with *Runx1* deletion alone or any other genotypes — developed AML (which we call primary or 1° AML) (Fig. 4B). Morphologically defined myeloblasts were abundant in the peripheral blood, bone marrow, and spleen, and they infiltrated multiple organs, including the liver (Figs 4C and S9A). The malignant cells expressed the c-Kit receptor but not mature lineage markers, including Mac1 (Fig. 4C). When the bone marrow or spleen cells were transplanted from the Case #1 mouse to sub-lethally irradiated mice, the recipients died with AML in three months (Fig. 4E).

All of the other ENU-treated Runx1^F/F^; *Mx1-Cre* mice, with or without the *U2af1*(S34F) mutation died prematurely with myeloid pathology: increased percentages of myeloid cells in the peripheral blood, enlarged spleen (Fig. S8B), and extramedullary hematopoiesis. Low-grade dysplasia in the erythroid and neutrophil lineages was occasionally observed in the peripheral blood and bone marrow of moribund URC mice, consistent with the multi-lineage dysplasia associated with *U2af1*(S34F). These results suggest that ENU-treated mice with deletions of *Runx1* develop lethal forms of myeloproliferative neoplasm (MPN), with occasional dysplastic features in the presence of *U2af1* (S34F).

We speculated that the animals with lethal forms of MPN but no frank AML might harbor malignant clones capable of producing myeloid leukemias if introduced into healthy recipient mice. To test this idea, spleen and/or bone marrow cells from moribund URC mice with MPN were transplanted to sub-lethally irradiated mice, and the recipients were monitored for up to one year after transplant for development of AML (Fig. 4A). None of the animals receiving transplanted cells from Runx1^F/F^; *Mx1-Cre* mice (n = 11) developed AML or other hematopoietic malignancies. However, all recipients of spleen and/or bone marrow cells from two of the eleven ENU-treated URC donors (Cases #2 and #3 in Figs 4B, 4D and S9) died with AML 12 or 42 weeks after transplant (Fig. 4E). Several hematopoietic compartments, including blood, bone marrow and spleen, as well as other tissues such as the liver, were filled with immature c-Kit^+^ hematopoietic precursors in the recipient mice, but not in the donor mice (Figs 4D, S9B, and S9C).

The c-Kit^+^ cells were GFP^+^ (data not shown), confirming that they were derived from the donor mice. Results from secondary transplants, using spleen cells from recipient mice, confirmed that the malignancies were transplantable (Fig. S9D and S9E). We refer to the AMLs derived from Cases #2 and 3 as late AMLs to distinguish them from the 1° AML observed in one ENU-treated URC mouse (Case #1).

In summary, we have observed a total of three cases of AML arising from the bone marrow of URC mice after ENU treatment (Fig. 4B). In contrast, AML did not appear in any of the ENU-treated mice with one gene alteration (i.e. only *U2af1*(S34F) or only deletion of exon 4 of *Runx1*); in recipients of bone marrow transplants from such animals; or in mice that did not receive ENU. These findings are consistent with the idea that initiation of AML in our cohort requires *U2af1* (S34F), *Runx1* deficiency, and treatment with ENU. However, due to the low incidence of AML (derived from three out of sixteen URC mice), a larger cohort would be necessary to establish the requirement of all of these three factors, including mutant U2af1, for leukemogenesis in a statistically significant fashion.

### Identification of probable leukemogenic, ENU-induced mutations by whole exome sequencing of DNA from Cases #1 - 3

We performed whole exome sequencing (WES) on DNA from GFP^+^Lin^-^c-Kit^+^ (LK) cells from the spleens of Cases #1, #2, and #3 mice to seek additional genetic changes caused by ENU. DNA from GFP^+^CD3ε^+^ or GFP^+^CD19^+^ lymphocytes from each of the three affected URC mice was used to determine a reference sequence. In each leukemic sample, we identified more than 1000 somatic variants by WES (Table S5). The most common variants were single nucleotide substitutions, among which G to A (or C to T) transitions, A to G (or T to C) transitions, and G to T (or C to A) transversions were the most frequent (Fig. S10A). These changes are consistent with previously reported effects of ENU (Table S6).

*Case #1:* By reviewing the variants for likely participants in leukemogenesis in primary AML cells from Case #1, we identified a splice donor site mutation in *Tet2* that is predicted to disrupt correct splicing, and a missense mutation in *Gata2* homologous to the GATA2(R362Q) mutation found in human AML (Fig. 5A – 5F).

**Figure 5.**
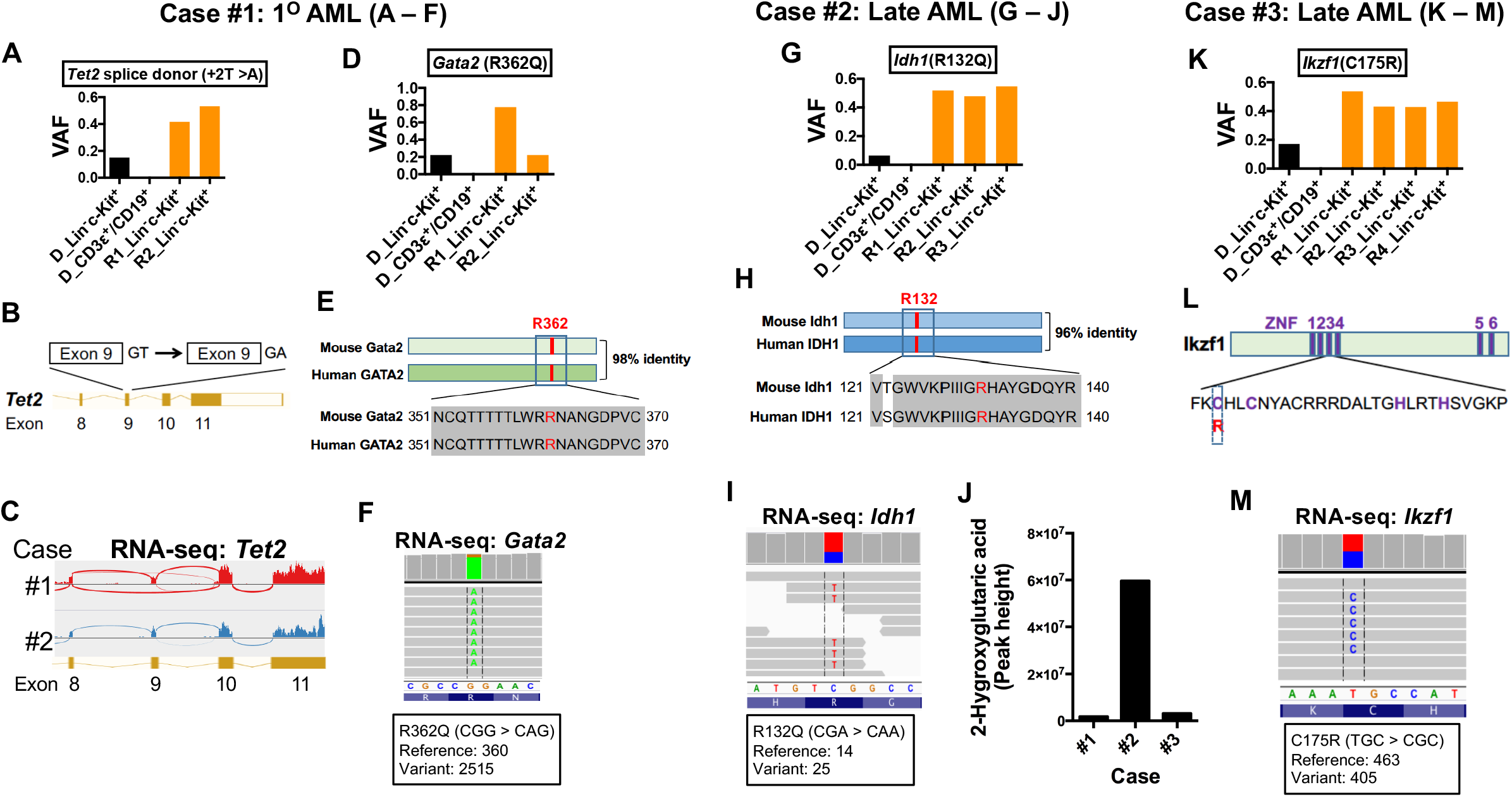
Acquired somatic mutations in Cases #1 – 3 AML cells affecting mouse homologs of human cancer genes. DNA was prepared from donor-derived (i.e. GFP^+^) Lin^-^c-Kit^+^ cells and lymphocytes (CD3ε^+^ or CD19+) from the spleens of donor and matched transplant recipient mice from Cases #1 – #3 and used for whole exome sequencing (WES) to identify somatic mutations. RNA-seq and metabolic analysis from some samples were performed to confirm the WES findings. (**A – F**) Acquired mutations in *Tet2* and *Gata2* in AML cells from Case #1. (**A** and **D**) VAFs for *Tet2* and *Gata2* mutations in the indicated cells from the donor (**D**) and transplant recipient (**R**) mice. (**B** and **C**) The T to A mutation in *Tet2* disrupts a splice donor site at the 3’ end of exon 9, producing transcripts in which exon 9 is omitted as shown by RNA-seq in the Sashimi plot in panel C. (**E**) Comparison of mouse Gata2 and human GATA2 proteins, highlighting in red the conserved arginine residue that was changed to glutamine residue by the missense mutation. (**F**) Confirmation of the *Gata2* (R362Q) mutation by mRNA sequencing from AML cells from a recipient mouse. (**G – J**) An Idh1(R132Q) mutation in AML cells from Case #2. (**G**) VAFs for the R132Q allele in DNA from donor and recipient mice. (**H**) Position of the mutated amino acid residue in mouse Idh1 and human IDH1 proteins. (**I**) Confirmation of Idh1(R132Q) by sequencing of RNA from AML from a recipient mouse. (**J**) AML cells with Idh1(R132Q) had elevated levels of 2-hydroxyglutarate, as measured by LC-MS/MS. (**K – M**) An Ikzf1(C175R) mutation in Case #3. (**K**) VAFs of the C175R allele in donor and recipient mice. (L) A cartoon illustrating the position of C175 in the third zinc finger domain of Ikzf1. (**M**) Confirmation of the *Ikzf1* (C175R) mutation by sequencing RNA from AML cells from a recipient mouse. Additional characterizations of the acquired mutations are presented in Fig. S10 and Tables S5 and S7.

*Case #2:* In the leukemic cells from Case #2, we found a missense point mutation in *Idh1* homologous to human DH1(R132Q) (Fig. 5G – 5J).

*Case #3:* In the leukemia cells from case #3, we found a missense mutation affecting a critical cysteine residue in a C2H2 Zinc finger domain of the transcription factor Ikzf1 (C175R, Fig. 5K – 5M).

In general, the variant allele frequencies (VAFs) of these potentially pathogenic variants were low (0.2 or less) in the donor LK cells, absent in the donor’s normal lymphocytes, and near or even above 0.5 in the LK cells recovered from the recipient mice that had developed AML (Fig. 5A, 5D, 5G, and 5K).

We next validated the presence of these variants and others by deep sequencing of several DNA samples from leukemic cells from Cases #1 – #3, using a focused panel of cancer-related genes (Table S7). This approach detected additional variants at low VAFs that are likely to be subclonal (Fig. S10B and S10C). We further confirmed the expression or functional consequences of these variants by mRNA sequencing. For example, the splice site mutation in *Tet2* resulted in exon skipping (Fig. 5C); the Gata2(R362Q), Idh1(R132Q) and *Ikzf1*(C175R) mutations were observed in mRNA (Fig. 5F, 5I, and 5M); Idh1(R132R)-expressing cells had high levels of 2-hydroxyglutarate (Fig. 5J). These identified variants are likely drivers of leukemogenesis, because *TET2* loss-of-function mutations, GATA2(R362Q), and *IDH1* (R132) mutations are often seen in human AML (see Discussion).

### The *U2af1*(S34F) allele is absent in AML cells derived from Cases #2 and #3

Encountering mutations in these mouse AMLs that are identical or similar to mutations frequently reported in human AML was surprising, but analysis of the DNA sequencing results revealed one additional surprise: *U2af1*(S34F) was absent in the DNA of LK cells from the transplant recipients from cases #2 and #3 that developed late AML (Fig. S10F and S10G, top panels). This was true despite the facts that Cre-mediated recombination was complete in the late AML cells and there was no evidence of focal deletions by PCR (Fig S10F and S10G, bottom panels; data not shown). The LK cells from these recipients were also GFP^+^ (data not shown) and showed complete deletion of the floxed exon 4 in the *Runx1* alleles (Fig. S10D), further confirming that these cells originated from the donors. The absence of *U2af1(S34F)* was also confirmed by the allele-sensitive Taqman assay, Sanger sequencing, and mRNA sequencing (Fig. S10H and data not shown). In contrast, *U2af1(S34F)* was present in LK cells from the donor Case #2 and #3 mice (albeit at a lower VAF), in matched lymphocytes and tails, and in the LK cells from Case #1 with primary AML (Figs S10E – S10G; data not shown). Moreover, the AML cells from Case #1, but not those from Cases #2 and #3, exhibited *U2af1*(S34F)-associated splicing changes in the consensus sequences preceding the altered cassette exons (data not shown), providing functional evidence for the presence or absence of *U2af1*(S34F) in these samples. These findings are considered further in the Discussion.

## DISCUSSION

Here we report the establishment of genetically modified mouse strains that express a commonly observed mutant allele, *U2af1*(S34F), from the endogenous *U2af1* locus in a Cre-dependent manner (Figs 1A and S2A). We show that production of *U2af1*(S34F) from one of the two *U2af1* loci in heterozygous (S34F/WT) mice alters pre-mRNA splicing in mouse cells in ways similar to those observed in *U2af1*(S34F)-expressing human cells, as previously reported by us and others (Figs 1B – 1E, S1D, S2E, and Table S3) (11, 21–24, 27). In the heterozygous mice, hematopoiesis is impaired, producing multi-lineage cytopenia and dysplasia (Figs 2, 3, and S4 – S6); these features are hallmarks of human MDS, a class of disorders in which mutant U2AF1, including *U2AF1* (S34F), or other mutant splicing factors are often found.

We also observed myeloid leukemogenesis in three of sixteen mice in which *U2af1*(S34F) is combined with homozygous deletion of *Runx1* and exposure to a mutagen, ENU, but not in mice lacking any of those three factors (Figs 4 and S9). Sequencing of DNA from the three AMLs identified additional mutations associated with human AML (Fig. 5; Table S5 and S7). However, the *U2af1*(S34F) missense mutation was absent in two of the three AML cases (Fig. S10F – S10H), suggesting that *U2af1*(S34F) may be involved in the initiation but not the maintenance of AML.

### U2af1(S34F) changes RNA processing in ways both common to and different from changes observed in human cells carrying *U2af1*(S34F)

Specific alterations in pre-mRNA splicing are the most widely characterized consequences of expressing *U2AF1* (S34F) in human cells (11, 21–24, 27). In this report, we show that *U2af1*(S34F) causes similar but not identical changes in mRNA splicing in both mouse fibroblasts and MP cells. Less than 5% of cassette exons show at least a 10% change in rates of inclusion in mouse cells expressing *U2af1*(S34F) (Figs 1E, S1D and S2E), numbers similar to those we previously observed in human cells expressing *U2af1*(S34F) (11, 21). Moreover, the consensus 3’ splice site sequences that precede the affected exons in mouse and human cells are very similar or identical. These similarities likely reflect the highly conserved sequence of U2AF1 protein and other components of the spliceosome machinery. For example, human and mouse U2AF1 proteins are nearly (more than 99%) identical, and the S34F mutations probably have similar effects on the two homologous proteins, recognizing highly similar 3’ splice sites in mouse and human transcriptomes.

Despite the similar effects of S34F mutations on sequence recognition and genres of altered splicing, the mouse mRNAs altered by *U2af1*(S34F) are not necessarily the homologs of the human mRNAs altered by *U2af1*(S34F) (Fig. 1B and 1D). Similar observations have been made previously in transgenic mice expressing *U2af1*(S34F) (27) and in mice expressing other mutant splicing factors (31–34). Despite many differences (roughly 20% of genes with mRNAs significantly affected by the S34F mutation in mouse and human cells are homologs), a few of the genes with altered mRNAs in both species are often mutated in human MDS and AML or are partners in gene fusions in myeloid disorders *(PICALM, GNAS, KDM6A* and *BCOR)* (35–38). Another gene *(RAC1)* plays an important role in HSCs (39, 40), and one more *(H2AFY)* encodes isoforms reported to affect erythroid and granulomonocytic differentiation (41). It remains to be determined, however, whether any of the changes in spliced mRNAs from these genes has a role in the hematologic abnormalities we have observed in our *U2af1(S34F)* mouse model.

Mutant U2AF1 has also been claimed to cause neoplasia by compromising mitophagy through alternative polyadenylation (APA) of *ATG7* mRNA (25) or by increasing the formation of R-loops (26). While we observed no evidence of effects of *U2af1*(S34F) on *Atg7* APA selection in mouse bone marrow cells (Fig. S1G), it remains to be determined whether R-loop formation is enhanced in mouse cells containing *U2af1*(S34F).

### U2af1(S34F) confers MDS-like features on mouse hematopoiesis but does not cause full-fledged MDS

We have found that expression of *U2af1*(S34F) from the endogenous locus in our mice impairs hematopoiesis, with features reminiscent of human MDS (Figs 2, 3, and S4 – S6). We also note similarities and differences in the hematopoietic phenotypes in our mice and the phenotypes in a previously reported transgenic mouse model (27), in which *U2af1*(S34F) is driven by a tetracycline-responsive element in the *Col1a1* locus. Cytopenia is a common feature of both models, but we observed cytopenia in all leukocyte lineages, whereas in the transgenic model, the affected cell populations are restricted to monocytes and B cells. Both models manifest defective HSC function since they compete poorly with co-transplanted normal HSCs in irradiated recipients. However, we observed decreased numbers of hematopoietic stem and progenitor cells (including LSK and HSC populations) in our mice, while the number of LSK cells was increased in the transgenic model. Furthermore, our mice developed mild macrocytic anemia and low-grade myelodysplasia, which were not present in the transgenic model. Factors contributing to these differences might include the levels of expression of the mutant proteins (mutant and WT proteins are at equal levels in our model but amounts may differ in the transgenic model) and the methods for inducing the mutant protein (transient treatment with poly (IC) to activate the *Mx1-Cre* transgene in our mice as opposed to addition of doxycycline to activate the rtTA transcriptional regulator in the transgenic model).

Mice engineered to produce mutant splicing factors other than mutant U2AF1 (31–34) display similar features, including cytopenia and a competitive disadvantage of mutant-containing HSCs in re-population assays. Therefore, different mutant splicing factors may elicit effects on hematopoiesis that are both common to splicing factor mutations and specific for certain factors.

Regardless of the hematopoietic phenotypes produced by different mutant splicing factors, none of the mouse models (including ours) recapitulates all aspects of human MDS. These results suggest that a mutant splicing factor is insufficient to drive human MDS on its own. Similarly, expression of an MDS-associated *SRSF2* mutant in human induced pluripotent stem cells did not affect the production of hematopoietic progenitor cells and only mildly affected hematopoiesis (42). In contrast, a much more drastic phenotype was produced by deletion of chromosome 7q, another common but much more extensive genetic alteration in MDS. These observations are consistent with findings that mutant *U2AF1* is often associated with other genetic alterations, such as deletion of chromosome 20q and mutations in *ASXL1, DNMT3A, TET2*, and *RUNX1* in human MDS (4, 6, 7, 12, 16, 17, 43).

Beyond MDS, the same mutations in *U2AF1* or other splicing factor genes are found in apparent healthy individuals with so-called Clonal Hematopoiesis of Indeterminate Potential (or CHIP) (44). HSCs from individuals with CHIP often carry loss-of-function mutations in *DNMT3A* and *TET2* (45, 46). The combination of *U2af1*(S34F) and loss of *Dnmt3A* or *Tet2* might produce hematopoietic phenotypes in mice that more precisely resemble human diseases. The occurrence of a *Tet2* splice site mutation in a case of AML in our mouse model (Fig. 5A – 5C) may reflect such cooperativity between mutant *U2af1* and loss of *Tet2.*

### Does *U2af1*(S34F) promote initiation of myeloid leukemia?

We have pursued the hypothesis that additional genetic changes are necessary to produce severe, progressive hematological disease in mice expressing *U2af1*(S34F) in the blood lineage. Although a larger cohort is necessary to reach statistical significance and confirm these findings, our results are consistent with the idea that *U2af1*(S34F) plays a necessary but not sufficient role in leukemogenesis in this setting. The results are also consistent with clinical observations that somatic mutation of *U2AF1* is a risk factor for AML in MDS patients (6, 12), in individuals with age-related clonal hematopoiesis (47), and in apparently healthy individuals (48).

We were surprised to find that in two cases of AML---the two detected after bone marrow transplantation—the leukemic cells contained only *U2af1* RNA with a WT coding sequence (Fig. S10F – S10H). These cells retained the Cre-recombined *MGS34F* allele (Fig. S10F and S10G) but codon 34 was composed of TCT (serine) rather than TTT (phenylalanine). A T-to-C transition might have occurred during exposure to ENU treatment or randomly during the leukemogenic process, and conferred a selective advantage of the kind demonstrated in the competitive transplantation experiments shown in Fig. 3. Regardless of the mechanism and significance of the loss of *U2af1*(S34F), our results suggest that *U2af1*(S34F) is dispensable for maintenance of murine AML in this setting. This is consistent with our previous observation that *U2af1*(S34F) is dispensable for the growth of established human lung adenocarcinoma cell lines carrying this mutation (11).

### ENU-induced mutations that likely cooperate with *U2af1*(S34F) and *Runx1* deletion in leukemogenesis

Exome sequencing of leukemic cells from the three cases of AML derived from our mouse model identified at least four somatic mutations that are likely to have contributed functionally to leukemogenesis (Fig. 5):

i. The *Tet2* variant in AML case #1 affects a splice donor site of a constitutive exon (exon 9, as shown in Fig. 5B) so that exon 9 is skipped during splicing (Fig. 5C). Loss of exon 9 (138 bp in length) does not change the reading frame in the Tet2 coding sequence, but loss of the 2-oxoglutarate-Fe(II)-dependent dioxygenase domain, partly encoded by exon 9, likely eliminates normal Tet2 function.
ii. The murine *Gata2* variant in case #1 AML is equivalent to GATA2(R362Q) (Fig. 5E), which is one of the recurrent *GATA2* mutations in AML [COSMIC ID: COSM87004; (49, 50)]. *GATA2* encodes a transcription factor that plays a critical role during hematopoiesis.
iii. The *Idh1* mutation identified in one of the late AML cases (Case #2) is equivalent to human IDH1(R132Q) (Fig. 5H). R132 mutants of IDH1, including IDH1(R132Q) produce the “onco-metabolite” 2-hydroxyglutarate (Fig. 5J) that is commonly found elevated in several human cancers including AML (51).
iv. The variant allele, Ikzf1(C175R), found in one case of late AML (Case #3), affects a conserved cysteine in a zinc finger domain of the transcription factor it encodes (Fig. 5L) and thus likely compromises (or alters) the DNA binding capacity of Ikzf1. *IKZF1*, a known tumor suppressor gene for lymphoid neoplasms (52), is also recurrently deleted in MPN (53) and pediatric AML (54). In combination with our results, it seems likely that *IKZF1* is a tumor suppressor gene for myeloid cancers as well.

These likely driver mutations from the three AML cases in our mice occurred largely independently, but some appear to be functionally related, implying that they may affect hematopoietic cell functions that must be altered to produce murine AML. For example, mutant IDH1 is known to inhibit TET2 enzyme activity (55), and Gata2 and Ikzf1 (as well as Runx1) are transcription factors critical for hematopoietic lineage specification and/or HSC function. Mutant U2AF1 might cooperate with alterations of the epigenetic modifiers and lineage transcription factors to initiate leukemogenesis.

## MATERIALS AND METHODS

### Generation of mice, MEFs, and related procedures

Details in generating the *U2af1*(S34F) conditional “knock-in” mice and related MEFs can be found in the Supplemental Information. All alleles, except for *Runx1^F^*, were used in a heterozygous (or hemizygous) state in this study. Mice carrying the *U2af1* targeted alleles were all backcrossed to B6 mice for at least three generations before they were used for the reported experiments, and were considered to have a mixed B6,129 background. Littermates of animals with experimental genotypes were used as controls in all described studies.

All procedures related to mouse were approved by the Institutional Animal Care and Use Committees at the National Human Genome Research Institute (NHGRI, Protocols: G-13-1 and G-95-8) and at Weill Cornell Medicine (WCM, Protocol: 2015-017). A brief description of these procedures can be found in the Supplemental Information.

### Cell culture, reagents, and assays

#### MEFs

Sub-confluent cells were treated with 4OHT (Sigma-Aldrich) at the indicated doses for 24 hr to induce Cre activity. The drug was then removed, and cells were cultured for an additional 48 hr before harvest. In other instances, the same treatment was repeated before harvest to improve the efficacy of Cre-mediated recombination.

#### Colony forming cell (CFC) assays

30,000 nucleated bone marrow cells were seeded in MethoCult GF M3434 medium (StemCell Technologies) in a 35 mm dish in triplicates, according to the vendor’s instruction. The number of colonies --- including erythroid (BFU-E), granulocyte-monocyte (CFU-GM) and multipotential (CFU-GEMM) colonies --- were counted 14 days later (or 7 days later when secondary plating was performed). When secondary plating was performed, primary CFC cultures from each set of triplicate cultures were harvested on day 8, pooled, and washed with IMDM containing 2% FBS. 30,000 or 10,000 nucleated cells were again seeded in a 35 mm dish with MethoCult GF M3434 medium in triplicate. The colonies were counted 12 days later. CFC assays was performed either as a contracted service (StemCell Technologies) or by the investigators.

#### RNA and RT-qPCR

RNA and RT-qPCR were performed as previously described (11). A brief description of these procedures can be found in the Supplemental Information.

#### Cell surface and intracellular staining and flow cytometry

Cell surface staining was conducted as previously described (56). RBCs in blood, bone marrow or spleen cell suspensions were lysed using ACK lysing buffer (Quality Biological). Description of the antibodies and procedures can be found in the Supplemental Information.

#### Metabolite quantification by LC-MS/MS

Two million live splenocytes from Case #1 – #3 recipient mice at the moribund stage were used to quantify for soluble metabolite by liquid chromatography coupled to tandem mass spectrometry (LC-MS/MS) at the Proteomics and Metabolics core lab (WCM) as a paid service.

### High throughput nucleotide sequencing

#### mRNA sequencing

mRNA sequencing (RNAseq) was conducted at either the Sequencing Facility (NCI) or the Genome Resources Core Facility (WCM). RNA quality was assessed by a Bioanalyzer (Agilent). Total RNA with good integrity values (RIN > 9.0) was used for poly A selection and library preparation using the Illumina TruSeq RNA library preparation kit. The MEFs samples were run on a HiSeq2500 instrument and sequenced to the depth of 100 million paired-end 101 bp reads per sample. All other samples were run on a HiSeq4000 instrument and sequenced to the depth of 50 million paired-end 51 bp reads per sample. These RNAseq data have been deposited into the NCBI GEO database (accession number GSE112174).

#### Whole exome sequencing (WES)

WES was performed in the WCM Genome Resources Core Facility. DNA quality was assessed with a Bioanalyzer (Agilent). A pre hybridization library was prepared using the KAPA LTP library preparation kit (Kapa Biosystems). Targets were enriched by NimbleGen SeqCap EZ Exome Library V2.0. Up to 18 samples were sequenced in one HiSeq4000 lane to generate paired-end 101 bp reads.

### Bioinformatics

Differential gene expression and splicing patterns were analyzed as described (11, 21). A brief description of these procedures can be found in the Supplemental Information.

For the analysis of whole exome sequencing data, raw sequencing reads were edited to remove low-quality bases, and adapter sequences were identified using cutadapt (57). The processed reads were then mapped to the mouse GRCm38 reference genome using BWA (58). The alignments were further processed for local realignment and base quality score recalibration using GATK (59). Somatic SNVs and small insertions and deletions (“Indels”) were detected using Varscan2 (60).

### Statistics

Statistical significance was determined by two-tailed Student’s t test using GraphPad Prism 6 or otherwise stated. In all analyses, p values ≤ 0.05 are considered statistically significant.

## ACKNOWLEDGMENTS

We thank Sukanya Goswami (WCM), Shao Ning Yang (WCM), Hayley Motowski (NHGRI), Jackie Idol (NHGRI), Danielle Miller-O’Mard (NHGRI), Ursula Harper (NHGRI), Amalia Dutra (NHGRI), Evgenia Pak (NHGRI), Stacie Anderson (NHGRI), Martha Kirby (NHGRI), David Bodine (NHGRI), Shelley Hoogstraten-Miller (NHGRI), Cecilia Rivas (NHGRI), and other members of the transgenic mouse core at NHGRI for exceptional technical assistance. We thank Nancy Speck (UPenn) for providing the conditional *Runx1* knockout mice. We thank Devin Koestler (KUMC) for consultation on statistical analysis. We thank members of the Varmus lab, the Liu lab, Guoan Zhang (WCM), Timothy A. Graubert (MGH), Pavankumar N.G. Reddy (MGH), Borja Saez (MGH), Matthew J. Walter (Wash.U.), Cara Lunn Shirai (Wash.U.), Dan Larson (NCI), Murali Palangat (NCI), and Markus Hafner (NIAMS) for helpful discussions during the course of the study.

HV and DLF were supported by the Intramural Program at the National Institutes of Health and are now supported by the Meyer Cancer Center at Weill Cornell Medicine and the Edward P. Evans Foundation. PL and TZ (Zhen) are supported by the Intramural Program at the National Institutes of Health.

## AUTHOR CONTRIBUTIONS

Conceived and designed the experiments: DLF, TZ (Zhen), PL, HV.

Performed the experiments: DLF, TZ (Zhen).

Analyzed the data: DLF, TZ (Zhen), BD, TZ (Zhang), RKB, PL, HV.

Contributed reagents/materials/analysis tools: JF, TZ (Zhang), LG, AY, OAW, RKB. Wrote the paper: DLF, HV.

---

**Supplemental Figure S1.**
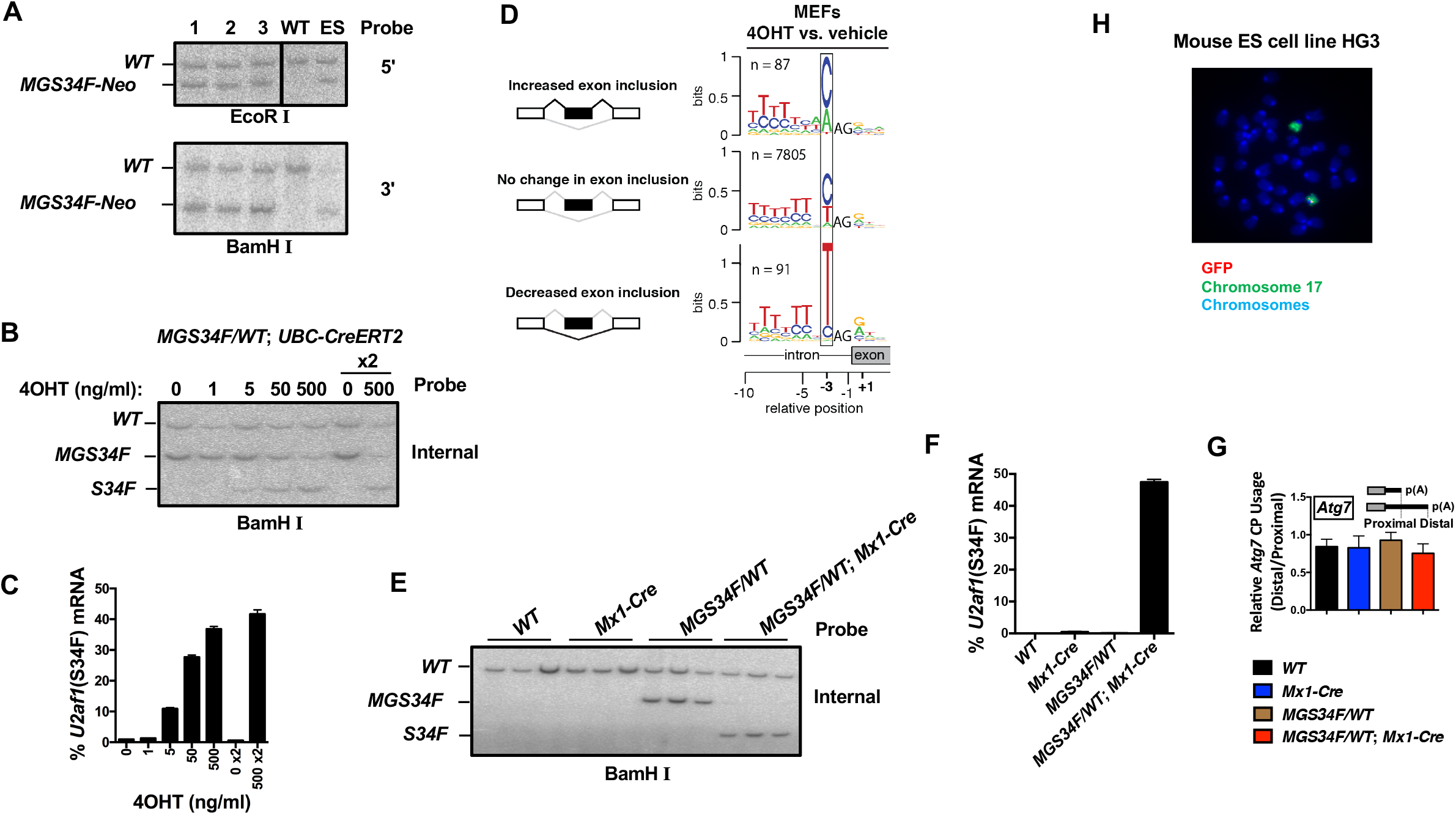

**Supplemental Figure S2.**
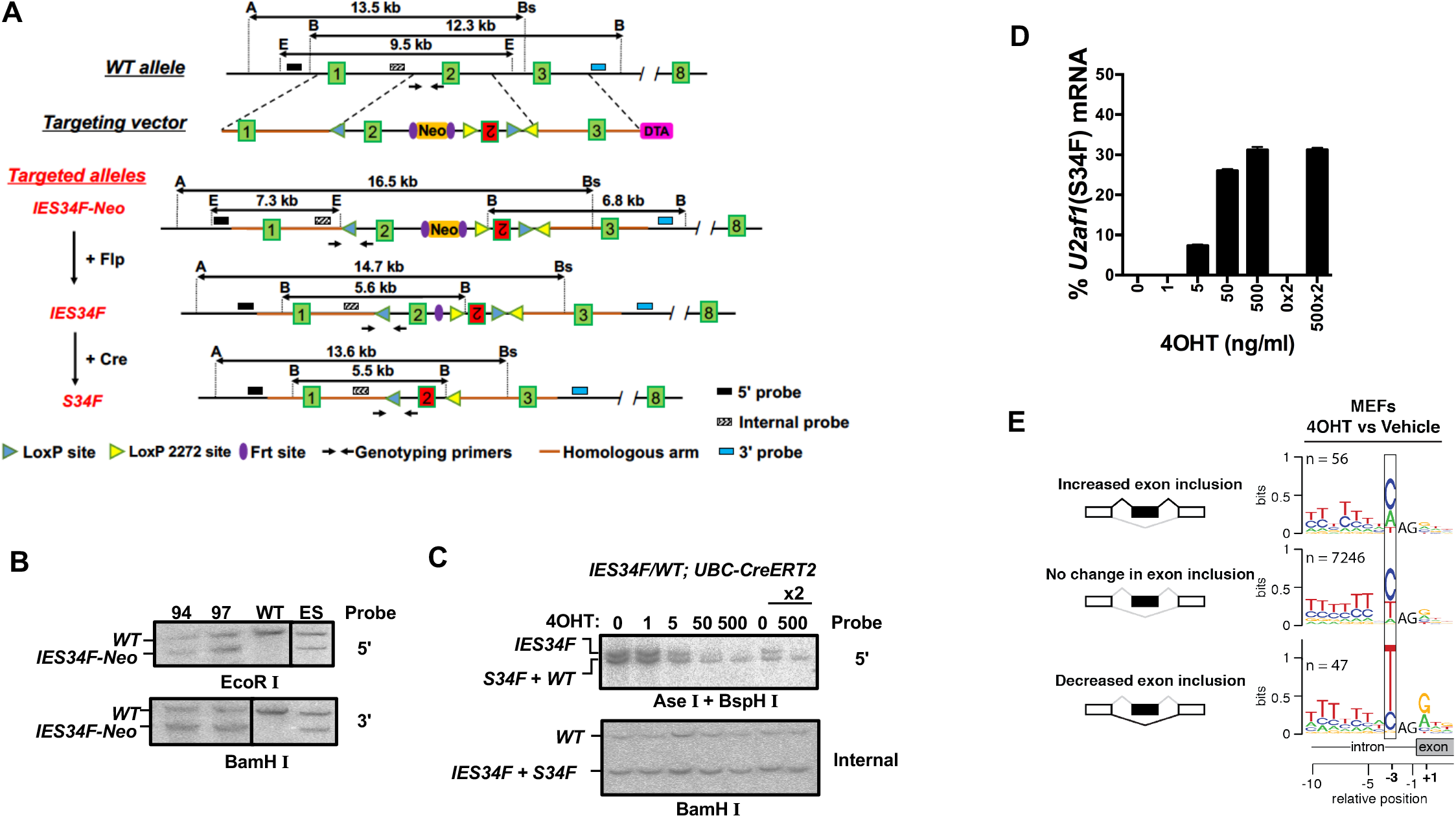

**Supplemental Figure S3.**
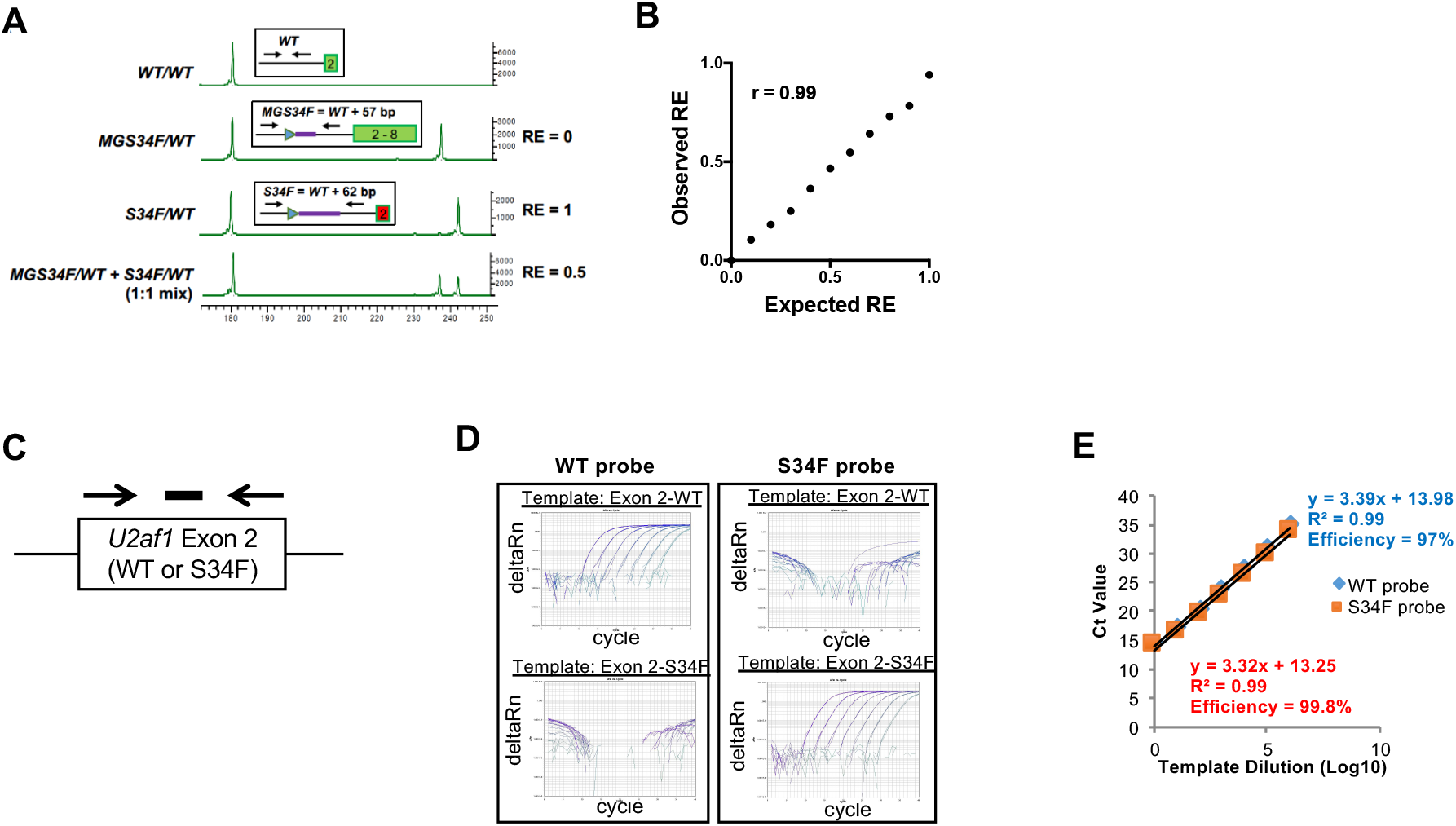

**Supplemental Figure S4.**
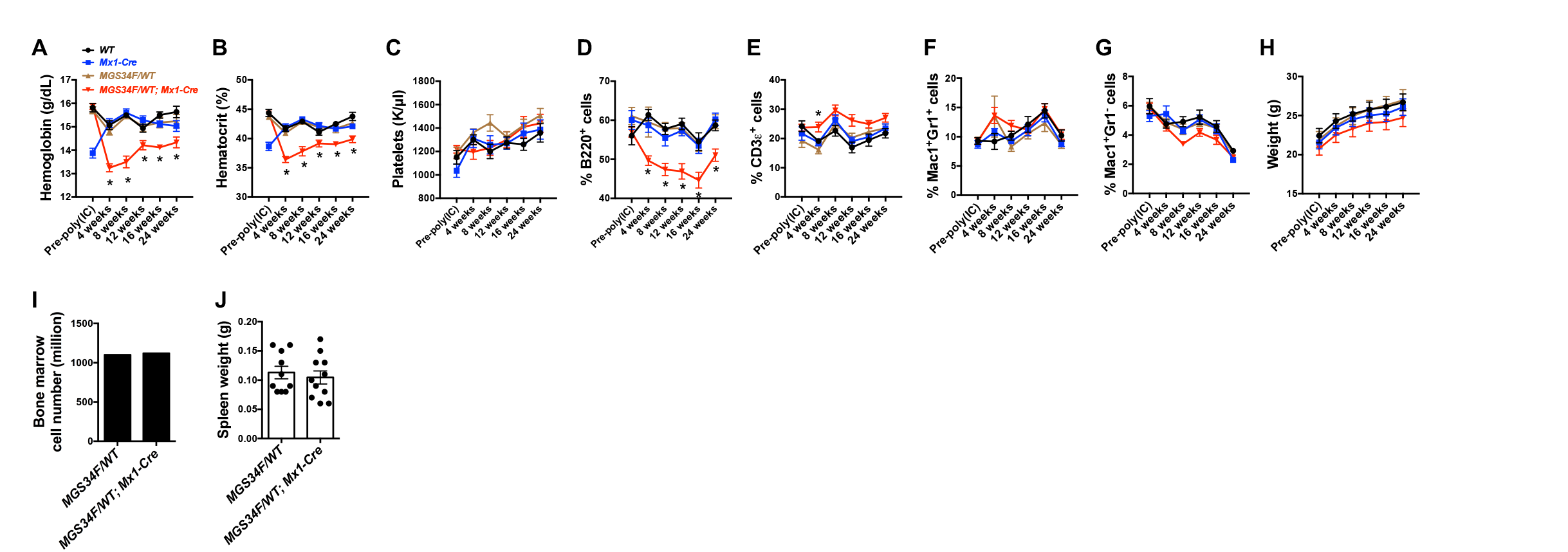

**Supplemental Figure S5.**
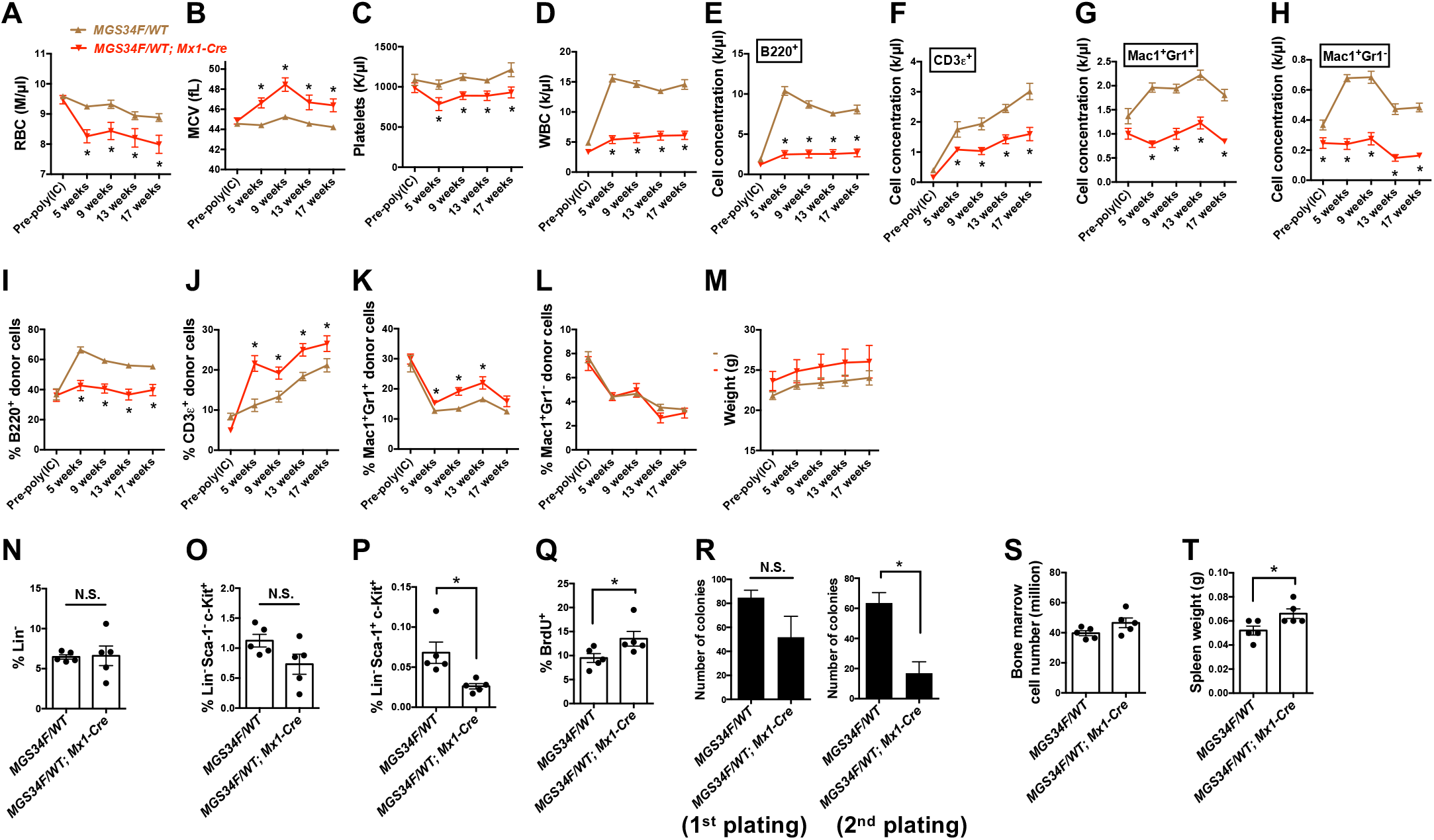

**Supplemental Figure S6.**
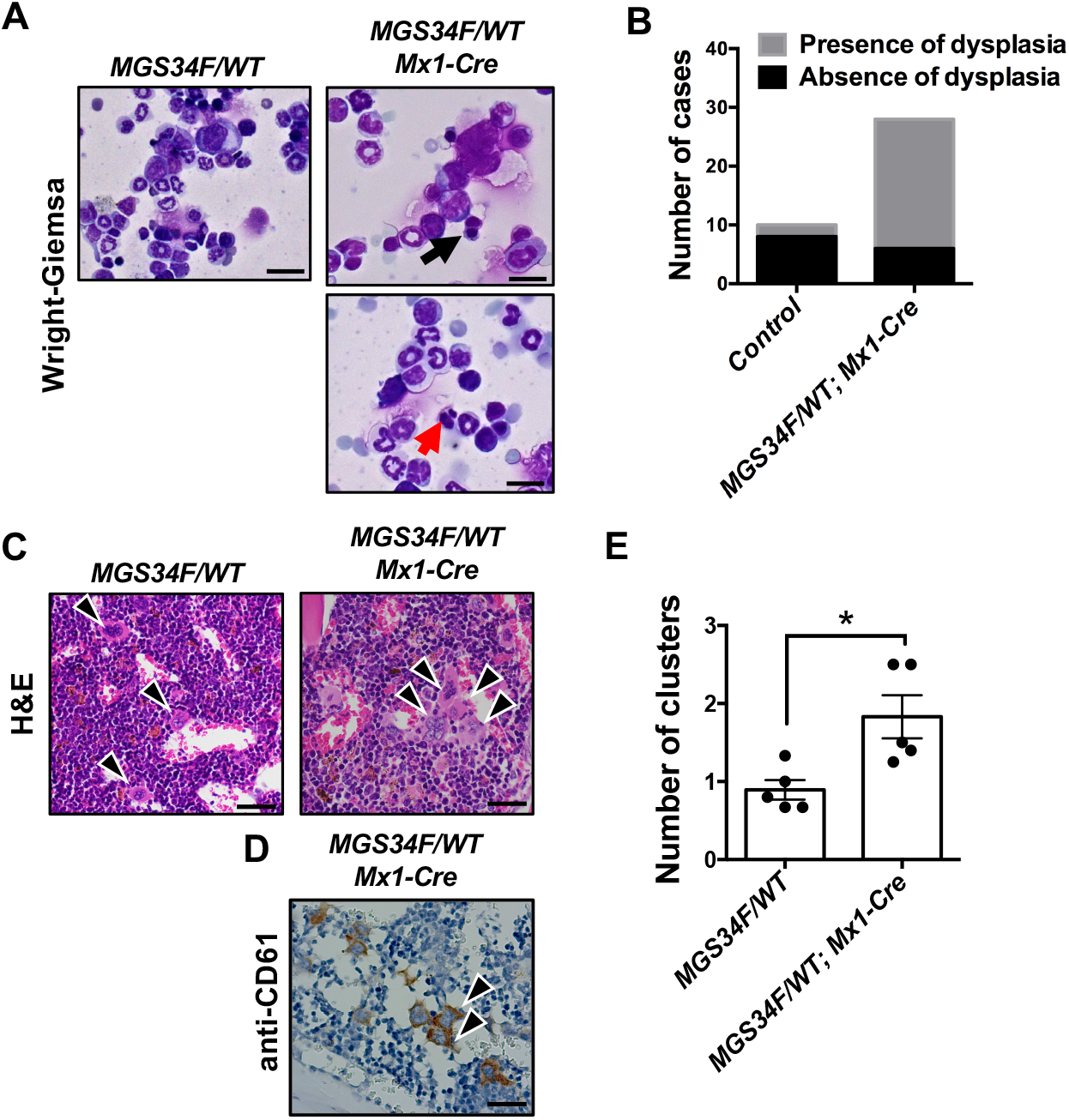

**Supplemental Figure S7.**
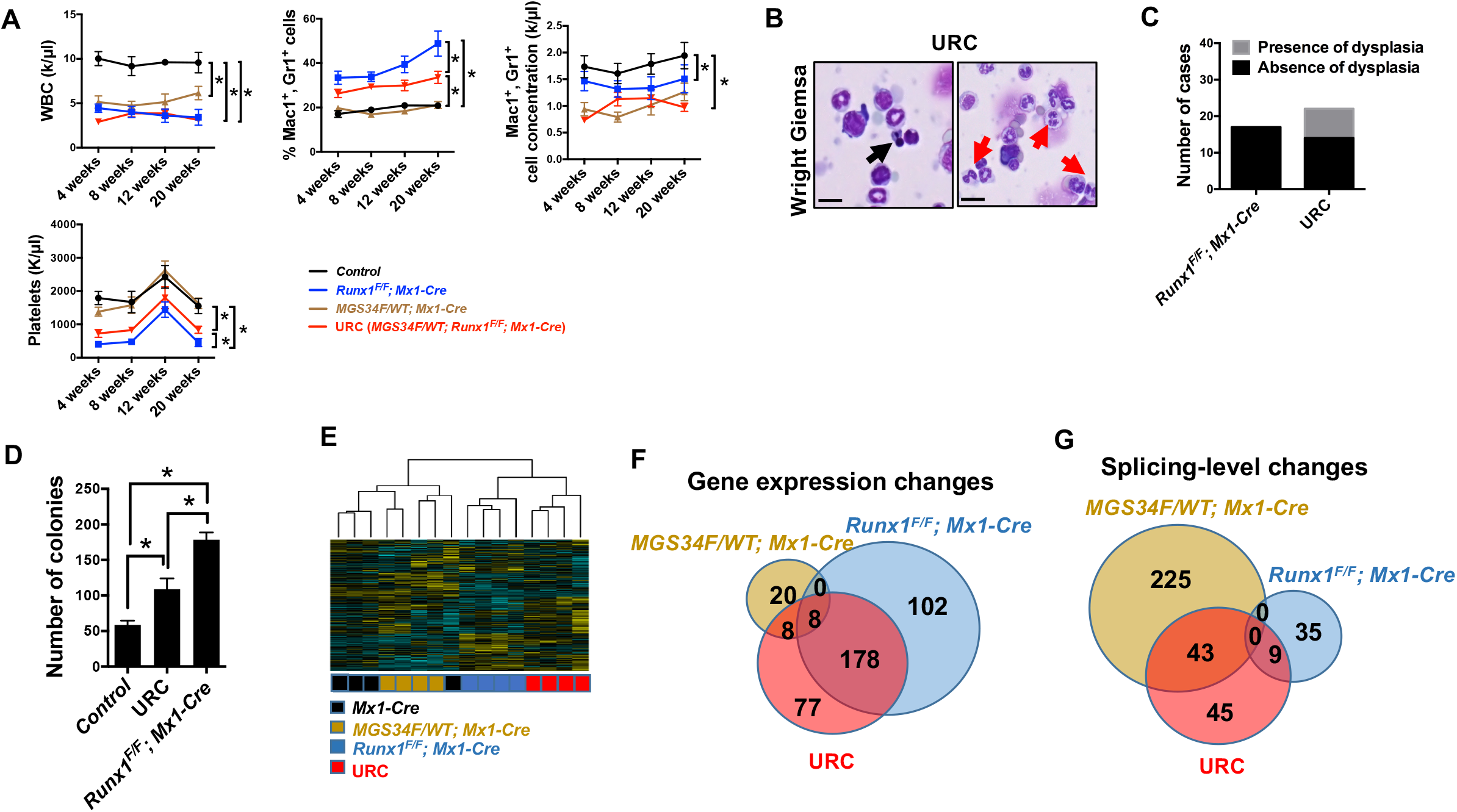

**Supplemental Figure S8.**
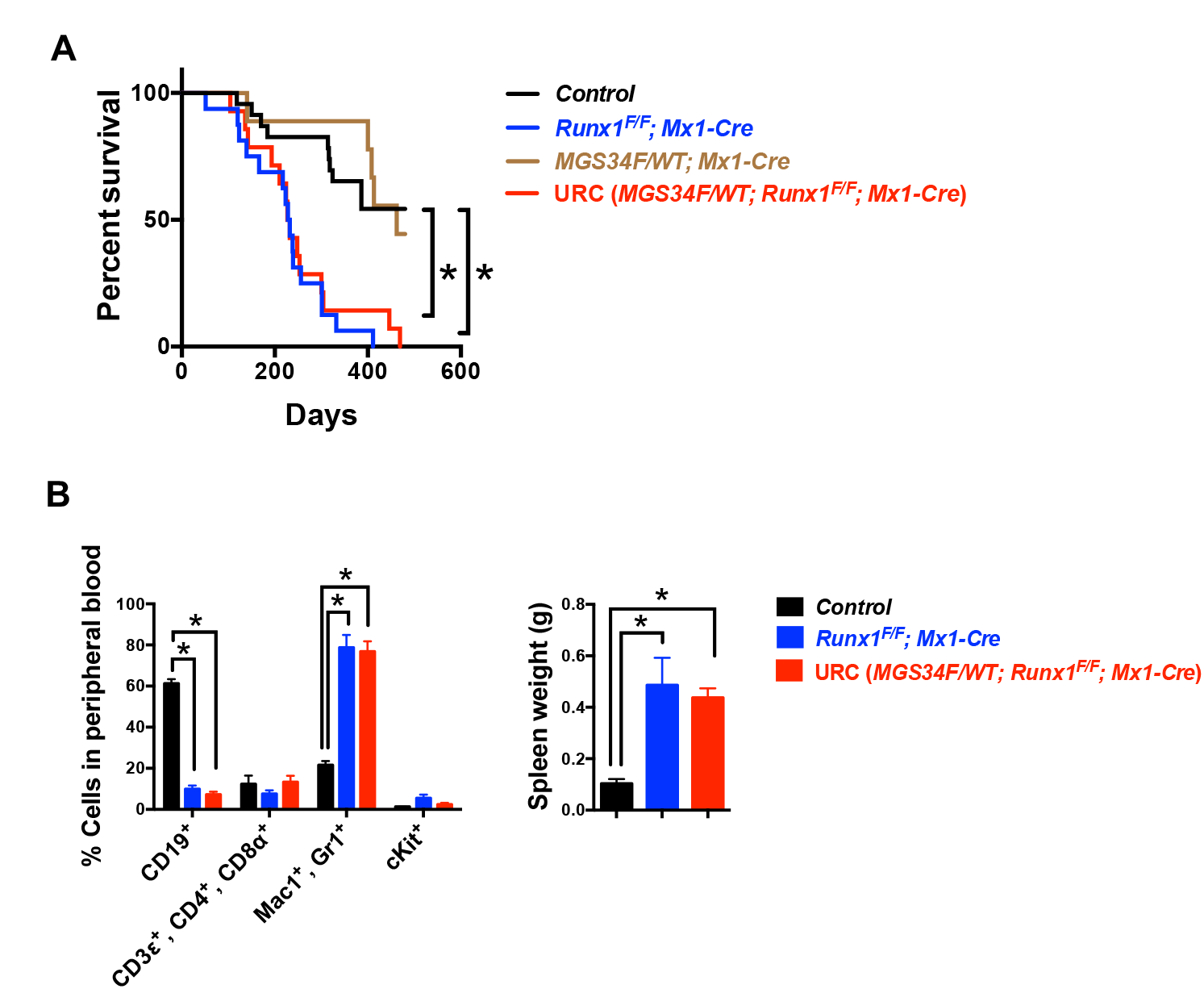

**Supplemental Figure S9.**
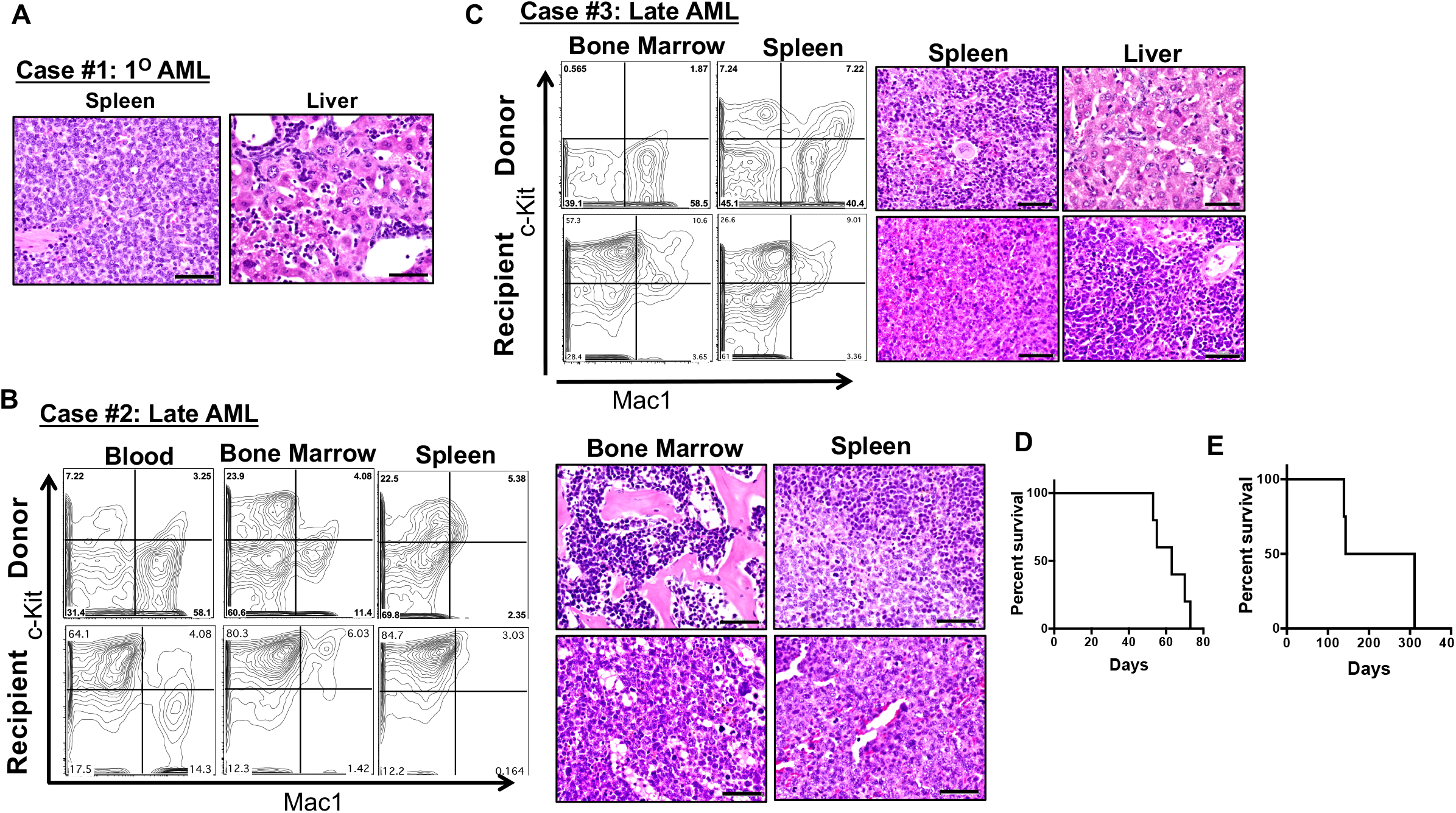

**Supplemental Figure S10.**
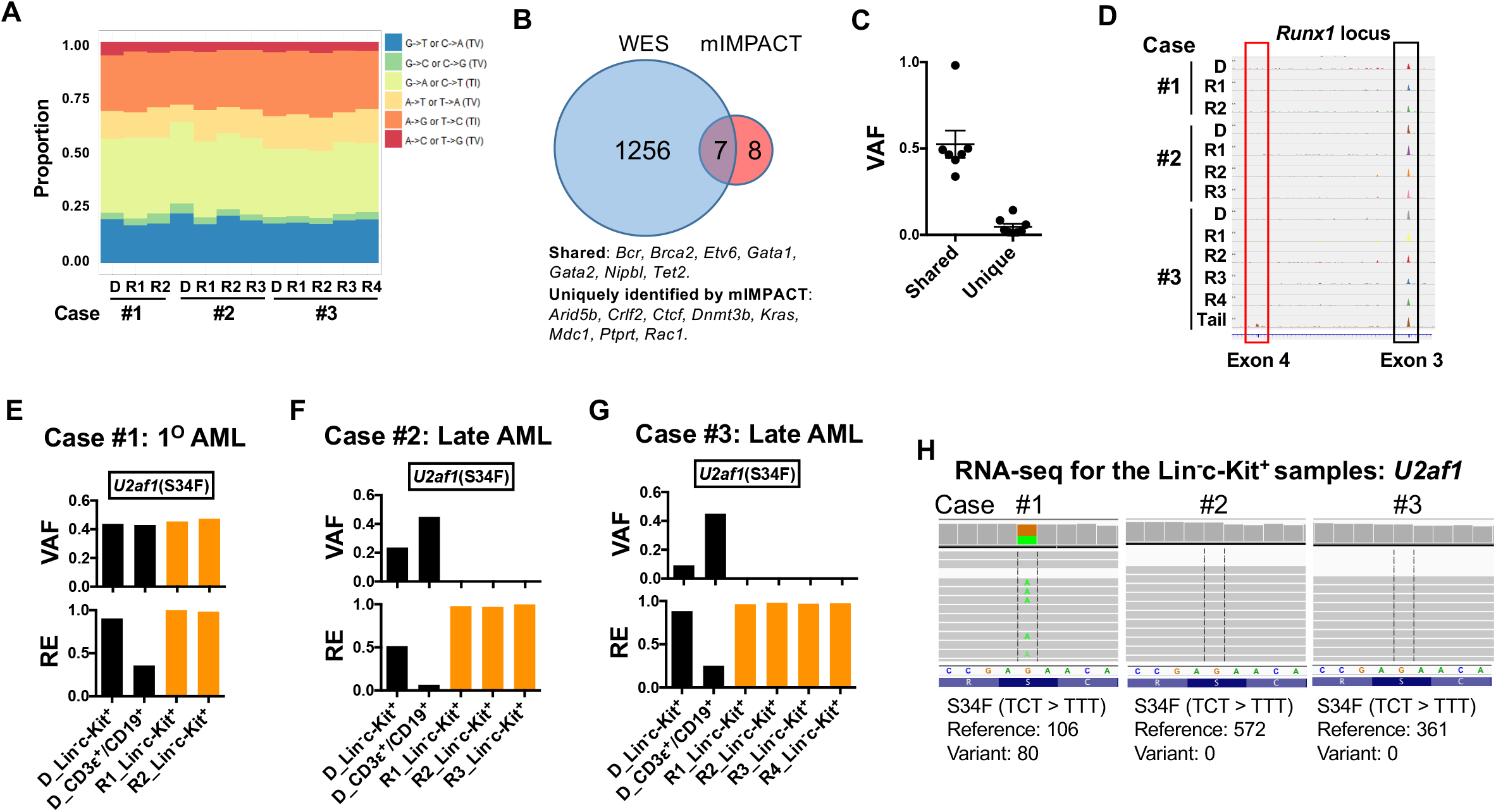

